# Spatial variation in exploited metapopulations obscures risk of collapse

**DOI:** 10.1101/315481

**Authors:** Daniel K Okamoto, Margot Hessing-Lewis, Jameal F Samhouri, Andrew O Shelton, Adrian Stier, Philip S Levin, Anne K Salomon

## Abstract

Unanticipated declines among exploited species have commonly occurred despite harvests that appeared sustainable prior to collapse. This is particularly true in the oceans where spatial scales of management are often mismatched with spatially complex metapopulations. We explore causes, consequences and potential solutions for spatial mismatches in harvested metapopulations in three ways. First, we generate novel theory illustrating when and how harvesting metapopulations increases spatial variability and in turn masks local scale volatility. Second, we illustrate why spatial variability in harvested metapopulations leads to negative consequences using an empirical example of a Pacific herring metapopulation. Finally, we construct a numerical management strategy evaluation model to identify and highlight potential solutions for mismatches in spatial scale and spatial variability. Our results highlight that spatial complexity can promote stability at large scales, however ignoring spatial complexity produces cryptic and negative consequences for people and animals that interact with resources at small scales. Harvesting metapopulations magnifies spatial variability, which creates discrepancies between regional and local trends while increasing risk of local population collapses. Such effects asymmetrically impact locally constrained fishers and predators, which are more exposed to risks of localized collapses. Importantly, we show that dynamically optimizing harvest can minimize local risk without sacrificing yield. Thus, multiple nested scales of management may be necessary to avoid cryptic collapses in metapopulations and the ensuing ecological, social and economic consequences.

## Introduction

Mismatches in spatial scale create pervasive problems in ecology and natural resource management (Cumming et al. 2006, Cope and Punt 2011). This problem occurs in part because the spatial extent of management or conservation units is often defined by history, jurisdictional, or institutional criteria rather than the scale of social and ecological processes at play (Levin 1992, Chesson 1998, Cumming et al. 2006). Such choices concerning the scale of management can result in spatial mismatches, where feedbacks controlling interactions among groups occur at different scales. In managed ecosystems like forestry and fisheries, mismatches may occur when harvest recommendations are based on trends in large-scale abundance without accounting for localized collapses (Johnson et al. 2012) or spatial variation in population structure and harvest rates (Cope and Punt 2011). Yet these spatially isolated collapses can have far-reaching consequences when the species play an indispensable role in local social-ecological systems, including human communities with limited capacity to forage over wide geographic scales. Empirical identification of appropriate spatial scales of management remains difficult for spatially structured populations, but can be a pre-requisite for diagnosing and reconciling challenges that spatial mismatches impose on the sustainable and equitable use of natural resources.

In metapopulations – plant or animal populations connected through dispersal – it is well established that the dynamics of individual populations can differ substantially from the aggregate metapopulation (Chesson 1998, Mangel and Levin 2005, Melbourne and Chesson 2006). A combination of movement, shared climate drivers, and compensatory processes determine whether dynamics of individual populations reflect the dynamics of the aggregate metapopulation (Kendall et al. 2000). Consideration of metapopulation structure has improved the management of the spotted owl (*Strix occidentalis*) (Lande 1988), salmon (*Onchorhyncus* spp.) (Stephenson 1999, Rieman and Dunham 2000, Schtickzelle and Quinn 2007, Peterson et al. 2014), amphibians (Marsh and Trenham 2001), and mosquitoes (Adams and Kapan 2009). To date, efforts to integrate metapopulation dynamics into natural resource management have largely focused on either minimizing the risk of localized extinction (Chadès et al. 2011), characterizing productivity of the metapopulation as a whole (sensu Takashina and Mougi 2015), or valuing benefits of portfolio effects (i.e. stabilizing effects of spatial asynchrony sensu Schindler et al. 2010). Less understood, both in theory and in practice, is if and when harvesting metapopulations can increase spatial variability that yield mismatches in spatial scales of management and population dynamics.

In this study, we assess how harvest dynamics interact with animal movement and recruitment to shape spatial population variability and risk of collapse at different spatial scales. Our results illustrate the challenge in managing spatially complex populations using three complementary approaches. First, we develop new theory using a stochastic analytical model to examine when and how harvesting in a metapopulation amplifies spatial variability that can create mismatches in spatial scale. Second, we contextualize the problem of spatial variability and mismatches in spatial scale by presenting historical analyses from spatial Pacific herring (*Clupea pallasii*) fisheries in British Columbia, Canada and home-ranges of associated predators and fishers that may be impacted by localized collapses. Finally, we evaluate how and when different harvest management approaches can ameliorate such mismatches using a numerical risk analysis in Pacific herring fisheries.

### Pacific herring case study in British Columbia’s Central Coast

Pacific herring exemplify the challenges inherent to managing metapopulations that exhibit spatial variation in population trends. In British Columbia (BC), Canada, herring return annually to nearshore coastlines in the late winter/early spring to reproduce. During this annual migration, they are harvested and preyed upon by a range of consumers. Mobile commercial fishing fleets harvest adult herring, largely for their roe, in the days prior to spawning. In contrast, Indigenous fishers are constrained to a local area and largely harvest eggs after spawning events (though some adults as well) as an important food, trade, and cultural resource (Lepofsky and Caldwell 2013, McKechnie et al. 2014, Department of Fisheries and Oceans 2015, von der Porten et al. 2016, Okamoto et al. 2019). These activities create trade-offs among commercial roe fisheries that remove spawning adults, which truncates adult age structure and reduces abundance, versus those that remove only eggs from shorelines (Shelton et al. 2014). Unfortunately, a core uncertainty for herring management, as for many species, is the extent of movement between areas (Flostrand et al. 2009, Benson et al. 2015, Jones et al. 2016, Levin et al. 2016). Similar uncertainty surrounds spatial variation in spawning biomass (Siple and Francis 2016), which may result in part from the degree of demographic synchrony between areas (e.g. synchrony in recruitment). Pacific herring in British Columbia are currently managed as stocks at regional scales (100s of kilometers) by Canada’s federal fisheries agency. Within these stocks, multiple spawning aggregations (substocks) seasonally occupy individual stretches of coastline, many of which are of important traditional and cultural value to Indigenous groups. Thus, Pacific herring fisheries present a valuable system in which to explore how spatial population dynamics, scales of management, and spatial constraints of fishers and predators interact to influence differential risk exposure to population collapses.

## Methods

### Methodological Overview

We used three distinct modeling approaches in this study to explore how harvest can affect spatial dynamics in metapopulations. In **Model 1,** we developed a novel analytical approach to modeling stochastic age-structured metapopulations to illustrate how and when harvest can interact with migration and both the magnitude and spatial synchrony of environmental stochasticity. We illustrated these important interactions using this model because of its simplicity and interpretability relative to more complicated nonlinear numerical models. In **Model 2,** we analyzed Pacific herring data from British Columbia using a Bayesian hierarchical model to estimate spatiotemporal variation in spawning biomass and harvest rates. In **Model 3,** to evaluate potential solutions for spatial mismatches and conditions for their success, we developed and applied a spatially explicit stochastic numerical model of metapopulations and their interaction with the fishery. Together, these approaches (Models 1-3) generate theoretical, empirical, and numerical results for understanding the complex, spatially-structured interactions among populations, harvest, and environmental variability. The collective results are then applied to evaluate the consequences for the availability of important natural resources for both human and non-human user groups.

### Theoretical effects of harvest on spatial variability in metapopulations (Model 1)

We used a simple metapopulation model to evaluate how harvest alters spatial variability, conditional on the underlying properties of the metapopulation. For our purposes, spatial variability is the degree to which volatility of the populations are masked by the observed volatility of the metapopulation.

We considered a simple metapopulation consisting of two populations, linked through the annual fraction migrating between populations (*δ*). We assumed both populations have the same dynamics (identical density independent adult total mortality rate (*Z*), maturity at age 2, Gompertz stock-recruit relationship, and symmetric adult annual migration) and trends in recruitment variability are controlled by a spatially correlated lognormal environmental stochasticity.

Let represent the abundance of age class *a* at location *i* at time *t as Y_a,i,t_*. Adult dynamics are shaped by mortality (*Z* - the sum of mortality from natural causes and harvest) and migration (0 ≤ *δ* ≤ 0.5):

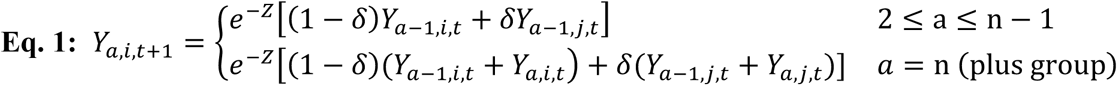

Total reproduction is the sum of adults across all adult age classes multiplied by 0.5 to account for an equal sex ratio:

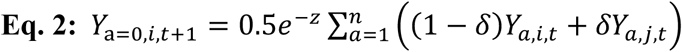

Both Eq. 1 and 2 are identical for both subpopulations (i.e. the system is symmetrical). We used a stochastic Gompertz model as the compensatory function that determines how zygotes translate into one-year-olds:

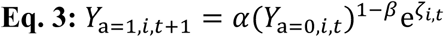

*α*, *β* and *ζ_i,t_* represent, respectively, the density independent productivity parameter, the within-location compensatory parameter, and environmental stochasticity that operates on post-dispersing larvae. The vector ***ξ_t_*** = [*ζ_1,t_, ζ_2,t_*]’ follows a multivariate normal distribution with mean zero and covariance controlled by the common within-site variance (σ_R_^2^) and spatial correlation (ρ_R_) yielding 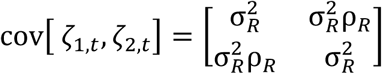.

To analyze the stochastic metapopulation model, we converted the model to a first order vector autoregressive model where statistical properties of stochastic forcing in multivariate systems are well described (Lütkepohl 2005). We first vectorized the model by aligning variables from both populations in a single vector:

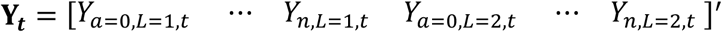

We then cast the model in terms of log-scale deviations from the equilibrium (i.e. 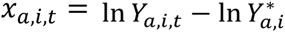 where 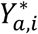 is the equilibrium) and linearized about the equilibrium (Nisbet and Gurney 1982, Bjørnstad et al. 2004) to create a first order vector autoregressive model. This approach approximates the deterministic dynamics of the nonlinear model with a Jacobian matrix (**J**) of first order dependencies and an environmental covariance matrix.

The matrix of Jacobian coefficients (with plus groups summed at age *n*) are partial derivatives for each age class within each subpopulation with respect to each other age class in each subpopulation. 𝐉 can be represented by the block matrix comprised of matrices describing among and within population age transitions:

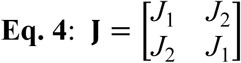

Where *J*_1_ is within population dynamics and *J*_2_ is among population dynamics defined by:

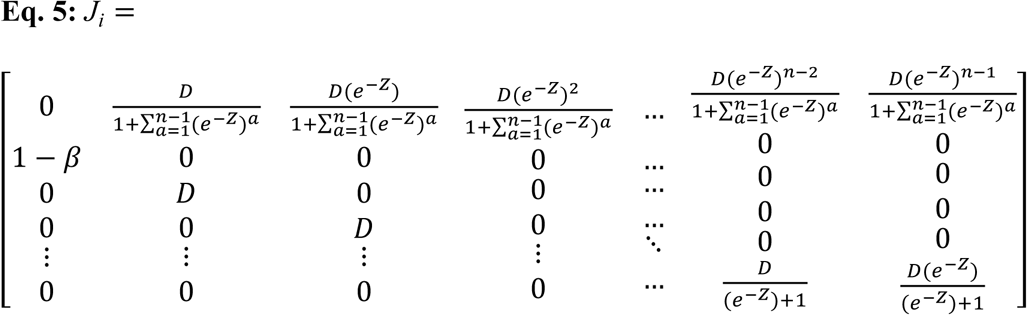

If *J_i_* = *J*_1_, *D* = (1 − *δ*) and if *J_i_* = *J*_2_, D = *δ*. *Z* is the annual total mortality rate and *n* is the number of age classes. See Appendix S1 for full derivation of the Jacobian and VAR(1) properties. The resulting VAR(1) model is:

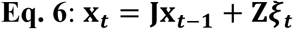

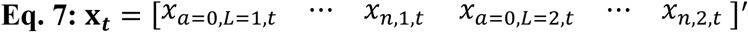

**Z** is a 2n x 2 binary matrix (2 age classes for each location, two locations) that controls which of the entries in **x***t* are subject to stochasticity from ***ξ****_t_* (i.e. translating the 2×1 vector of location specific stochasticity to a 2n x 1 sparse vector). In this case, only age class 1 (corresponding to the second column of *J_1_*) for each subpopulation is subject to stochasticity.

We used the known statistical properties of a VAR(1) model (Lütkepohl 2005) in tandem with the moments of a multivariate lognormal to derive the coefficient of variation for the subpopulations and metapopulation that yields the spatial variation in the metapopulation. Finally, to evaluate how local environmental sensitivity of population growth changes with mortality rate, we used a first-order impulse response analysis (Lütkepohl 2005) to estimate the annual intrinsic growth rate response to a recruitment pulse. The impulse response in our case describes how a recruitment perturbation affects the intrinsic growth rate of each subpopulation. Specifically, it is given by the entry of the 2^nd^ column of the 1^st^ row of the Jacobian matrix in Eq. 5. See Appendix S1 for full derivations and details.

### Pacific herring case-study: spatial variability in biomass and catch (Model 2)

To estimate patterns of spatial and temporal variability in herring biomass and harvest rates, we used spawn deposition and harvest data for six major Pacific Herring spawning units that comprise the Central Coast stock in British Columbia, Canada. Briefly, SCUBA surveys are used to estimate herring egg abundance which is converted to total spawning biomass with the assumed conversion of ∼100 eggs/g and an equal sex ratio (DFO 2015). For consistency we use the same index as in the DFO stock assessment but in the spatially disaggregated form. Catch (which occurs in the days prior to spawning) is reported by geographic section that are delineated and aggregated geographically. For full description of the time series see DFO (2015) and references therein.

To estimate spatial biomass trends from the survey and harvest time series observations, we used a nonlinear model in a Bayesian hierarchical state-space framework. We assumed survival, reproduction, and competition were location specific. We modeled expected change in biomass in year t+1 using a combined growth and survival model (individual growth and survival from year t) and Gompertz recruitment model (from year t-1, because fish mature at or after age 2 - Table 1, Eq. 8).

**Table 1:** Equations used in Model 2 (Pacific herring case-study: spatial variability in biomass and catch)

| Eq | Description | Equation |
| --- | --- | --- |
| Eq. 8 | Expected pre-spawn biomass | $\hat{b}_{t+1,l}^+ = \exp(\alpha_l + (1 - \beta_l) \ln(b_{t-1,l})) + \lambda_l b_{t,l}$ |
| Eq. 9 | Estimated log-biomass | $\ln \mathbf{b}_t^+ = \ln(e^{\ln \mathbf{b}_t} + \mathbf{h}_t) \sim \text{MVN}(\ln \hat{\mathbf{b}}_t^+, \text{diag}(\boldsymbol{\sigma})\Omega)$ |
| Eq. 10 | Objective function | $\ell = \sum_{t=1}^T \sum_{l=1}^L \frac{(\ln(\text{obs}[b_{t,l}]) - (\ln b_{t,l} + \ln q))^2}{(2\sigma_{obs}^2)} - \ln(\vert d\vert )$ |
| Eq. 11 | Jacobian adjustment for change of variables in Eq. 10 | $d = \frac{\partial}{\partial \ln b_{t,l}} \ln(e^{\ln b_{t,l}} + h_{t,l}) = \frac{b_{t,l}}{b_{t,l} + h_{t,l}}$ |
| Symbol | Description | Prior or Data Source |
| $\alpha_l$ | Site-specific Gompertz productivity | $[\alpha_l \quad \text{logit}(\beta_l) \quad \text{logit}(\lambda_l)]' \sim \text{MVN}(\boldsymbol{\Theta}, \boldsymbol{\sigma}_{\Theta}\Omega_{\Theta});$ |
| $\beta_l$ | Site-specific Gompertz compensation | |
| $\lambda_l$ | Site-specific aggregate mortality & somatic growth | |
| $\boldsymbol{\Theta}$ | Mean vector for $\alpha$, $\text{logit}(\beta)$, and $\text{logit}(\lambda)$ | $\boldsymbol{\Theta} \sim \text{MVN}(0, \text{diag}(1.5)\Omega_{\bar{\Theta}});$ |
| $\sigma_{\Theta_l}$ | Among site variances in $\alpha$, $\text{logit}(\beta)$, and $\text{logit}(\lambda)$ | $\sigma_{\Theta_l} \sim \text{half-cauchy}(0, 2.5)$ |
| $\Omega$ | Process error spatial corr. matrix | LKJ prior (Lewandowski et al. 2009): scale = 2 |
| $\Omega_{\Theta}$ | parameter corr. matrix | |
| $\Omega_{\bar{\Theta}}$ | mean parameter corr. matrix | |
| $\boldsymbol{\sigma}$ | Vector of site-specific lognormal process variances | $\sigma_l \sim \text{normal}(\sigma_{\mu}, 0.15); \sigma_{\mu} \sim \text{normal}(0, 0.15)$ |
| $\sigma_{obs}$ | Lognormal obs. error | $\sigma_{obs} \sim \text{half-cauchy}(0, 2.5)$ |
| $\ln q$ | Survey bias (egg “catchability”) | $\ln q \sim \text{normal}(0, 0.05)$ (Martell et al. 2012) |
| $\text{obs}[b_{t,l}]$ | Input Data | observed spawning biomass at time t at location l |
| $h_{t,l}$ | Input Data | harvest at time t at location l |

We estimated both process error and observation error in the model. We estimated spatially correlated process error that represents deviations from the expected log-scale pre-harvest biomass within each location. Process error may arise from a diverse combination of factors including immigration or emigration and temporal variation in growth, recruitment, or mortality (Table 1, Eq. 9). We also estimated location-specific observation error (Objective function - Table 1, Eq. 10), and a common survey bias (a “catchability” coefficient quantifying the mean proportion of eggs that are observed in fishery-independent surveys - Table 1, Eq. 10). Thus, we estimated trends conditional on estimated survival, biomass growth, process error covariance, observation error, and survey bias.

Because we estimated log-scale pre-harvest biomass but observations are of post-harvest biomass and harvested biomass, the model requires a change of variables and thus the posterior requires a Jacobian adjustment of the inverse transform (Gelman et al. 2013, Carpenter et al. 2016, Table 1, Eqs 10, 11). See Table 1 for all model equations, parameter definitions, and prior distributions. For details of estimation, posteriors, and model validation, see Appendix S2. We estimated the models using Stan (Stan Development Team 2016b, a) with 3 independent Markov Chains with 1000 iteration chains after 1000 iteration burn-in. We confirmed chain mixing and convergence using Gelman-Rubin statistic (R<1.01, Gelman and Rubin 1992) and for mdel adequacy and model fit using posterior predictive checks (see below). Posteriors compared with priors for core parameters are shown in Appendix S2: Fig. S1.

We used the full model posterior of post-harvest biomass to calculate the following metrics: 1) temporal and spatial variation in biomass (see Appendix S2: Fig. S1), and 2) the exploitation rate. We compared the estimated local harvest rates from the model posterior to the theoretical proportional allocation where fishing mortality is constant in space and an optimized allocation of harvest given the quota (see Appendix S2: Fig. S3 for results, Appendix S3 for methods) and the posterior mean. Here, we defined “optimized allocation” of catch as one that distributes catches in space according to the ideal free distribution (see Appendix S3 for methods). This distribution removes proportionally more biomass from subpopulations with higher biomass and is the spatial allocation of catch that minimizes effort to achieve the overall quota (assuming catch per unit effort is linearly related to biomass).

### Solutions for spatial scale mismatches in fished herring metapopulations (Model 3)

We used a stochastic model to simulate how spatial metapopulation dynamics and alternative management scenarios interact to influence risk of collapse at the subpopulation and metapopulation scales under a diverse suite of scenarios. The scenarios we examined include a factorial gradient of 1) annual movement (see Annual adult migration among spawning areas below), 2) environmental recruitment synchrony (the degree of correlation in recruitment in space – see Spatiotemporal Recruitment Dynamics below), 3) a range of harvest rates (see Stock Harvest Quota below), and 4) allocation of harvest in space (see Spatial Harvest Prosecution below). For this analysis, we defined a “collapse” as years with spawning biomass below 20% of unfished biomass. We used this definition for herring because, 1) it is generally seen as a conservative estimate of biomass below which traditional Indigenous harvest becomes difficult and, 2) it lies below the current closure threshold of 25% unfished biomass. Alternative thresholds defining collapse were also assessed but yielded qualitatively similar results (data not shown).

For all analyses, we simulated 10 populations placed around a hypothetical circular island of arbitrary size, where distances among adjacent populations were equal. Fish spawning at any site in a given year were able to move to any other site in the next year. The probability of migration from one site to another declines as alongshore distance between the locations increases and is controlled by a periodic kernel. Likewise, synchrony in stochastic recruitment among sites decays with distance between locations controlled by a periodic covariance kernel. Stochasticity in dynamics was included in recruitment (spatial and temporal variation), survival (temporal variation only) and movement probabilities (spatial and temporal variance). The order of operations mathematically was 1) recruitment, 2) survival, 3) movement, 4) roe fishery harvest, and 5) spawning. Details of the simulation are outlined below with core equations listed in Table 2 and definitions in Table 3.

**Table 2:** Equations used in Model 3 (Solutions for spatial scale mismatches in fished herring metapopulations). See Table 3 for parameter values and definitions.

| Eq | Description | Equation |
| --- | --- | --- |
| Eq. 12 | Recruits (age 2) prior to movement | $n_{a=2,l,t+2}^{++} = \text{eggs}_{l,t} \exp \left[ \epsilon_{l,t} - \frac{\sigma_r^2}{2} \right] / (\alpha + \beta \text{eggs}_{l,t})$ |
| Eq. 13 | Total eggs laid | $\text{eggs}_{l,t} = \sum_{a=2}^{10} m_a f_a [0.5 n_{a,l,t}]$ |
| Eq. 14 | Environmental recruitment stochasticity | $\epsilon_t = \phi \epsilon_t + \text{MVN}(0, \sigma_r^2 [1 - \phi^2] \Omega_r)$ |
| Eq. 15 | Spatial correlation in recruitment stochasticity | $\rho_{i,j} = \exp \left( - \frac{2 \sin^2 \pi \vert \text{dist}_{i \rightarrow j} \vert }{\eta^2} \right)$ |
| Eq. 16 | Pre-harvest abundance | $n_{a,l,t+1}^{++} = \begin{cases} \lambda_t n_{a-1,l,t} & 2 < a < 10 \\ \lambda_t n_{a-1,l,t} + \lambda_t n_{a,l,t} & a = 10 \text{ (plus group)} \end{cases}$ |
| Eq. 17 | Post-migration abundance | $N_t^+ = N_t^{++} S; S_{i,j} = P(i \rightarrow j)$ |
| Eq. 18 | Recruit or Adult migration by distance function | $P(i \rightarrow j) = \exp \left( - \frac{2 \sin^2 \pi \vert \text{dist}_{i \rightarrow j} \vert }{\gamma^2} \right) \sum_{j=1}^N \exp \left( - \frac{2 \sin^2 \pi \vert \text{dist}_{i \rightarrow j} \vert }{\gamma^2} \right)^{-1}$ |
| Eq. 19 | Quota | $\hat{Q}_{t+l} = \begin{cases} \min(H_{\text{Target}} * \hat{B}_{\text{forecast},t}, \hat{B}_{\text{forecast},t} - L_{\text{crit}} * \hat{B}_{0,t}) & \text{if } \hat{B}_{\text{forecast},t} \geq L_{\text{crit}} * \hat{B}_{0,t} \\ 0 & \text{otherwise} \end{cases}$ |
| Eq. 20 | Post-harvest abundance at age in year t | $n_{a,l,t} = (1 - m_a) n_{a,l,t}^+ - m_a n_{a,l,t}^+ e^{-F_{t,l}}; F_{t,l} = \ln h_{t,l} / \sum_{a=2}^{10} w_a m_a n_{a,l,t}^+$ |
| Eq. 21 | Biomass at location l in year t | $B_{l,t} = \sum_{a=2}^{10} w_a m_a n_{a,l,t}^+$ |

**Table 3:**
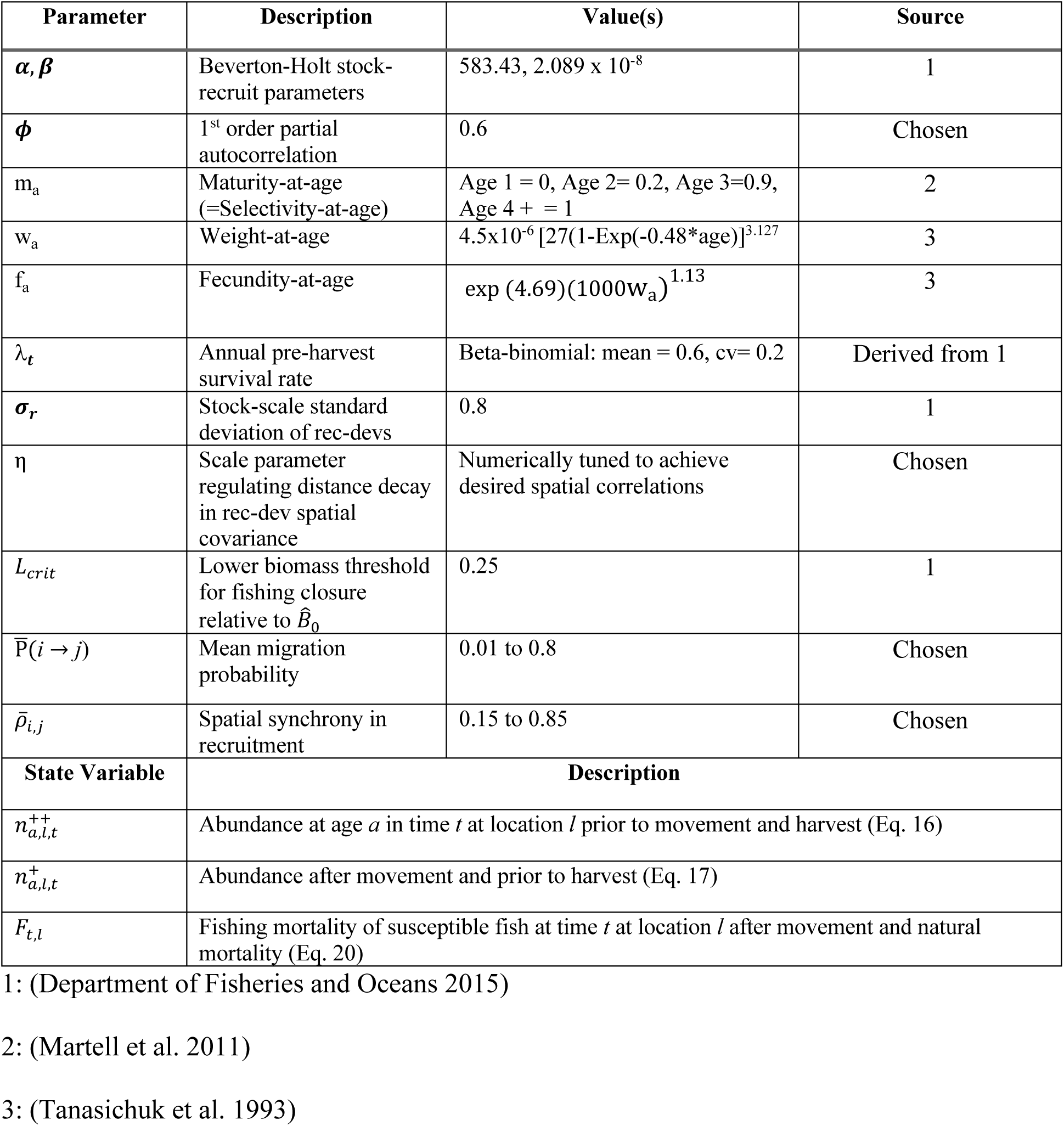
List of key parameters and state-variables in Model 3 with definitions, values and citations. Note that equilibrium considerations are generated by simulating with no harvest and no stochasticity (*σ_r_* = 0, cv in *λ_t_* = 0, and harvest = 0). Selectivity-at-age in all cases is equal to maturity-at-age.

#### Spatiotemporal Recruitment Dynamics

Following the assumptions of the current British Columbia herring assessments (DFO 2015), recruitment of age 2 individuals at location i is a function of eggs produced 2-years prior and local density dependence via a Beverton-Holt model. We allowed recruitment to exhibit both spatial and temporal variability and autocorrelation with a first-order vector autoregressive model. Spatial correlations in recruitment followed a Gaussian spatial decay with distance (d), and location specific recruitment variability was tuned such that net recruitment variability was approximately constant across scenarios (CV of metapopulation recruitment = 0.8; DFO 2015).

#### Annual Adult Survival and Migration Among Spawning Areas

All adult survival occurred prior to movement into spawning locations, was constant in space and was randomly drawn from a beta-binomial with mean λ *=* 0.6 with coefficient of variation of 0.2 (DFO 2015). Survivors migrated to new spawning locations with a probability that decays with distance from the previous site, controlled by a periodic kernel, tuned to achieve the desired retention rate.

#### Stock Harvest Quota

We followed the existing harvest control rules of Pacific herring in British Columbia (DFO 2015). The annual biomass harvest quota for the stock 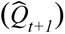 was designed to achieve a target harvest rate (H_target_) and a minimum spawning escapement (25% of B_0_ – the average steady state biomass without fishing). For simplicity and to avoid evaluating stock assessment model performance (which is out of the scope of this study) we assume forecasts are unbiased with no observation noise.

#### Spatial Harvest Prosecution

We used and compared two spatial fleet allocation scenarios to generate the distribution of spatial commercial roe harvest given the stock quota: 1) the fleet prosecuted the fishery equally in space whereby the absolute harvest was directly proportional to spawning biomass within a given year (***proportional allocation***) or 2) the fleet was allocated according to the ideal free distribution (IFD) that in theory would optimize catch efficiency if fleets are not spatially constrained (***optimized allocation -*** See Appendix S3 for details in generating the IFD). Here, realized catch was nonlinearly related to spawning biomass, harvesting more from areas with higher biomass and leaving alone areas with lower biomass. See Appendix S3 for methods and results from a third allocation strategy, a random spatial allocation that is more similar to the fleet allocation in the empirical case study. In all cases, fishery selectivity was identical to maturity-at-age reflecting that harvest occurs on mature fish at the spawning grounds.

#### Simulation details

For each simulation, we 1) evaluated deterministic equilibria without fishing, 2) initiated stochastic forcing of recruitment from the equilibrium for 12 years (allowing the full suite of age classes to be influenced by environmental stochasticity), 3) initiated the fishery in year 13, and 4) recorded performance metrics for years 23-52. We ran 100 replicate simulations for each combination of migration probability and recruitment synchrony. Primary performance metrics summarized for each simulation included a) mean number of years below 20% B_0_ at both stock and substock scales to assess risk of collapse, b) mean stock and substock level catch, c) mean stock and substock biomass, d) stock and substock temporal variability (coefficient of variation) in biomass, and e) mean spatial variation in biomass (difference between squared coefficient of variation at the substock versus stock scale).

## Results

### Theoretical effects of harvest on spatial variability in metapopulations (Model 1)

Our model illustrates that spatial variation among subpopulations increases with higher harvest rates (Fig. 1a) and decreases with synchronizing forces of migration and environmental correlation (difference in surfaces in Fig. 1b). As a result, the metapopulation trend and coefficient of variation are less reflective of trends and variation in its subpopulations as harvest increases (Fig. 1b). Biologically, this higher mortality rate decreases longevity and increases local-scale sensitivity to spatially explicit recruitment pulses. The amplification of fluctuations occurs at a higher rate at local scales than on aggregate. This response decreases spatial coupling and predictability. Reduced longevity of adults (via higher mortality) reduces the abundance of adults in each subpopulation that buffers against local stochasticity through survival and migration.

**Figure 1:**
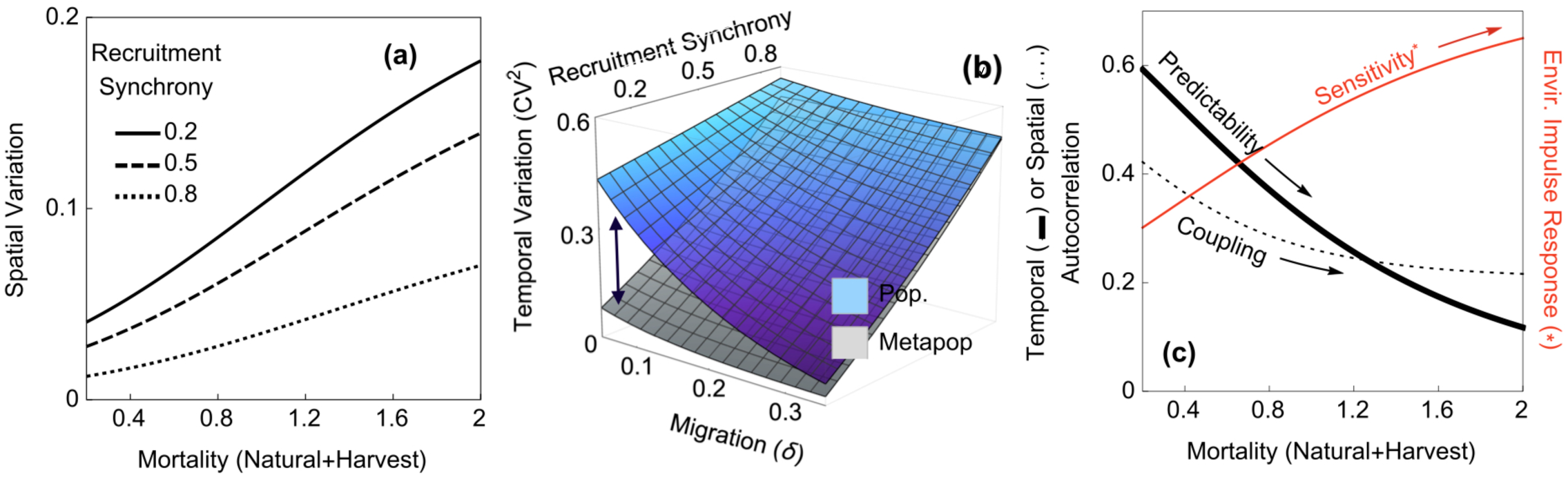
Effect of elevating harvest mortality rate on asynchrony related spatial variability in the metapopulation from Model 1. **(a)** Spatial variation increases as a function of total mortality for different levels of spatial synchrony in recruitment productivity. Here migration is held at 0.25 and log-scale recruitment variability (σ_R_) is 0.7 (i.e. CV = 0.8). Note spatial variation is a result of the discrepancy in population versus metapopulation temporal variation in panel (b); **(b)** Temporal variability (squared coefficient of variation) of the population and metapopulation. The arrows illustrate spatial variability as the difference between surfaces. **(c)** Measures of local population predictability (measured by first order within population autocorrelation, thick black line), among population coupling (measured by among population correlations, thin black line), and local environmental sensitivity (measured by the response at the population level to a unit environmental impulse affecting recruitment productivity, red line).

Mathematically, this result emerges for several related reasons. Increases in mortality (constant across space in this case) reduce local subpopulation inertia (predictability, measured as 1^st^ order temporal autocorrelation, thick black line in Fig. 1c). This reduction in inertia increases subpopulation sensitivity to temporal variation in local subpopulation recruitment (red line in Fig. 1c). This sensitivity is illustrated by the primary impulse response at a single lag which in this case always increases with total mortality (Z) derived from Eq. 5:

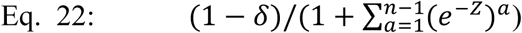

Such increases in local environmental sensitivity reduce spatial coupling (measured as the spatial autocorrelation, dotted black line in Fig. 1c) and thereby increase spatial variability in abundance.

Importantly these patterns of spatial variability can persist even in the presence of modest migration rates (Fig. 1b). While spatial variability decreases with synchronizing forces of migration (δ) and environmental correlations (ρ_R_) (Fig. 1b, see Appendix S1 for solutions), spatial variability only diminishes as migration and environmental correlations are substantially high (i.e. as migration probabilities approach 0.5 to produce 100% mixing). This phenomenon is illustrated by the discrepancies between temporal variability of the metapopulation and component subpopulations that produce spatial variation (Fig. 1b).

Next we illustrate challenges imposed by spatial variation in exploited metapopulations using an empirical case study of Pacific herring and present solutions using a numerical management strategy evaluation.

### Pacific herring case-study: spatial variability in biomass and catch (Model 2)

Pacific herring subpopulations on the Central Coast of BC exhibit substantial spatial variability in subpopulation trends (Fig. 2a, b, c). The estimated biomass of individual (“local”) subpopulations has varied by more than an order of magnitude over the past three decades, and similar differences in biomass are evident among subpopulations in the same year. As a consequence, aggregate (“regional”) stock biomass at any one point in time is bolstered by a few subpopulations, while others linger at low levels (Fig. 2b, Appendix S2: Fig. S2). Synchrony in biomass among subpopulations is low (0.29), with much higher variability at the subpopulations scale (average CV = 0.88) than in aggregate (CV = 0.54). This discrepancy results from high spatial variability in subpopulations trends (Fig. 1c - also known as *β* variability; (Wang and Loreau 2014). In fact as much as an estimated 91% of an individual subpopulation’s biomass is harvested annually, though the aggregate exploitation rate fluctuates around the target of 20% (Fig. 2d, e, Appendix S2: Fig. S3). Counterproductively, this can result in occurrences where subpopulations experiencing periods of lower-than-average biomass are heavily exploited preceding collapse (e.g. Fig. 2d, in 2006). Harvest rates generally differ in magnitude among subpopulations in any given year (Appendix S2: Fig. S3) and higher harvest rarely focus on the subpopulations with highest biomass (Appendix S2: Fig. S3). This is illustrated directly by spatial harvest distributions that deviate substantially from baselines of spatial harvest evenness used here and in the simulation model (proportional vs optimized allocation - Appendix S2: Fig. S3). Next we explore the potential consequences of different spatial harvest distributions via a closed-loop simulation model.

**Figure 2:**
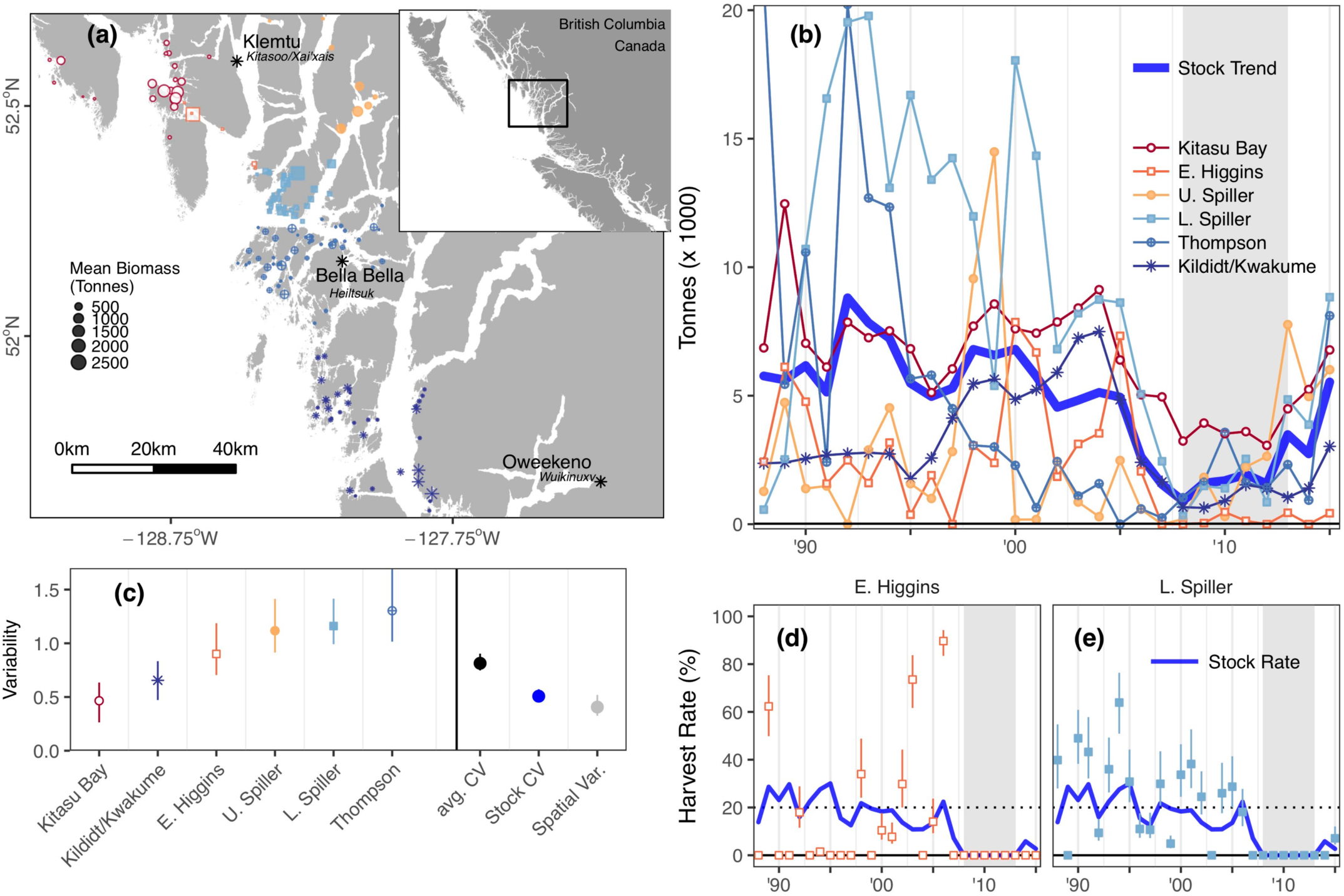
(a) Map of major Pacific Herring spawning substocks (“Sections” - denoted by different symbols) as defined by Canada’s Federal Fisheries Agency in the Central Coast of British Columbia. Indigenous communities noted with *. **(b)** Trends for the six main supopulations (substocks - thin colored lines & points) and the metapopulation (stock) mean (thick blue line). Trends were estimated using multivariate hierarchical Bayesian state-space model integrating spawn surveys and catch information (model 2). Individual plots for each substock are shown, along with catch, spawn observations, and 95% posterior credible intervals in Appendix S2. **(c)** Temporal coefficients of variation estimated from the Bayesian state-space mode over 25 years for each substock, the mean of each substock CV (large black symbol), the regional stock coefficient of variation (blue point), and the spatial variability (black point with error bar) which is the standardized variance of the system after accounting for the variance of the stock and the variance of the substock means. Error bars are 95% credibility intervals. (d) Trends in annual harvest rate for 2 of the 6 local substocks (Higgins and Lower Spiller) with points, 95% credibility intervals and the mean for the aggregate stock (thick blue line). For full results see Appendix S2. The dotted line represents the target harvest rate of 20%.

### Solutions for spatial scale mismatches in fished herring metapopulations (Model 3)

We used numerical simulation of a metapopulation to determine whether and under what conditions exploitation rates that appear sustainable in aggregate can risk collapse of local subpopulations, and by extension, adversely affect predators and the fishers who target them at this scale. The divergence in risk among scales, an effect of the mismatch in spatial scale, is controlled by the magnitude of harvest rates, allocation of harvest in space, annual migration, and spatial recruitment synchrony. Our simulations show that risk of collapse can be 10 times greater at local subpopulation scales than at aggregate metapopulation scales (Fig. 3) for the 20% harvest rule. While it may seem intuitive that relatively modest adult migration would minimize differences in risk to subpopulations and the metapopulation, our results do not support this supposition. Even relatively high migration rates can impose substantial discrepancies in the risk of collapse between subpopulation and metapopulation scales (Fig. 3). This principle holds so long as spatial synchrony in recruitment is not exceedingly high (Fig. 3 upper portions of heatmaps). For most scenarios, high spatial variability (i.e. from low annual migration among subpopulations or low spatial recruitment synchrony) leads to low apparent risk at the aggregate metapopulation scale, despite high risk of collapse for local subpopulations (Fig. 3; risk increases towards the lower left quadrants).

**Figure 3:**
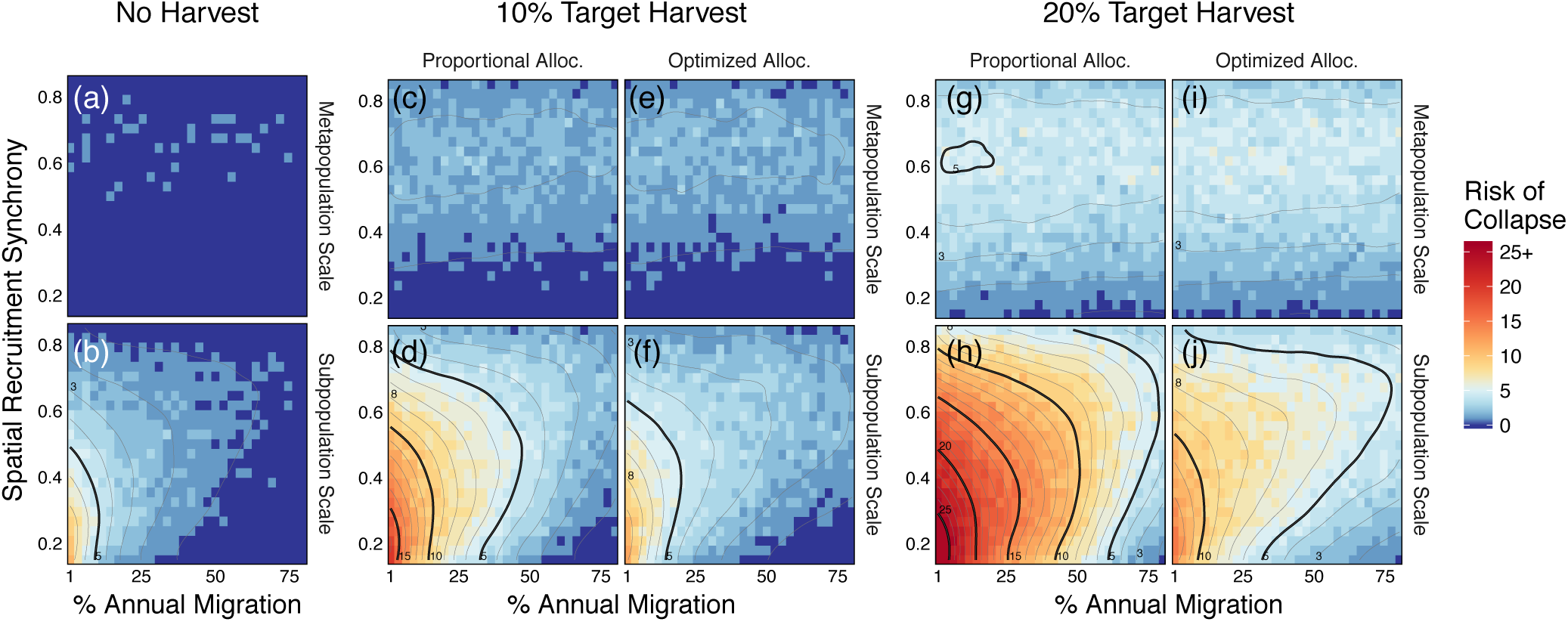
Mean risk of collapse (%) at metapopulation and subpopulation scales for different harvest rates and spatial harvest allocation strategies under varying spatial scenarios. Results are derived from simulation-based risk analysis of hypothetical Pacific herring fisheries. Axes present simulations with different levels of annual migration and spatial synchrony in recruitment productivity, columns represent different harvest strategies and rows represent the spatial scales of inference. Risk of collapse is the probability of falling below 20% of unfished equilibrium (i.e. the dotted lines in Fig. 4 a-c); at the subpopulation scale, risk is measured as the mean probability of collapse for each population. *Optimized allocation* harvests proportionally more from subpopulations with higher biomass; *Proportional allocation* removes the same proportion of biomass from each subpopulation. Annual migration is the mean percent of each subpopulation that emigrates each year and spatial synchrony is the mean pairwise correlation in recruitment productivity. For full results see Appendix S3: Fig. S2. Note that the allocation strategies produce equivalent average yields (Figure 3c vs. 3e and 3g vs. 3i; see Appendix S3: Fig. S3 for yield comparisons). For an illustration of how alternative strategies affect the mean duration of collapses, see Appendix S3: Fig. S4.

We find that risk to local subpopulations greatly exceeds risk to the aggregate metapopulation with both a simulated 10% and 20% target harvest rate. Under the 20% target harvest, risk of collapse at the local scale exceeds 10%, even with annual migration among subpopulations approaching 50% (Fig. 3 h). When harvest is optimized in space or target harvest rates are reduced to 10%, risks of collapse at local scales are substantially reduced (Fig. 3 d,f,j versus Fig. 3 h) and limited to scenarios where migration is ∼10-15% and spatial synchrony in recruitment is low. Local scale risks of collapse are even worse under a random (or opportunistic) spatial allocation, but also ameliorated by reductions in harvest (Appendix S3: Fig. S2). Importantly, a 10% target harvest with optimized spatial allocation never exceeded 10% risk of collapse in our simulations.

The discrepancy in risk of collapse at local subpopulation versus aggregate metapopulation scales is shaped by both overall harvest rates and spatial variance in subpopulation trend (Fig. S5). Risk at the subpopulation scale matches that of metapopulation scales when subpopulations exhibit little spatial variance (i.e. when there is high recruitment synchrony or high migration, top vs bottom panels in Fig. 3 c-j, SI Fig. S6). In contrast, risk diverges with increases in spatial variance (Fig. S5). Spatial variance is shaped not only by spatial recruitment synchrony and migration, but also harvest rates and allocation in space. Fig. 4b-c illustrates how different harvest allocations impact spatial variation in trends (Fig. 4 d, e). Higher harvest rates increase risk at subpopulation scales in part because of higher depletion at the metapopulation scale (Fig. 3 d,f,h,i) and also because harvest can increase spatial variance (Fig. 4d, also explored more generally in Fig. 1) when efforts are allocated proportionally. This result demonstrates how low demographic synchrony creates a portfolio effect at the metapopulation scale but simultaneously masks risks of collapse at the subpopulation scale which can be exacerbated with higher harvest rates.

**Figure 4:**
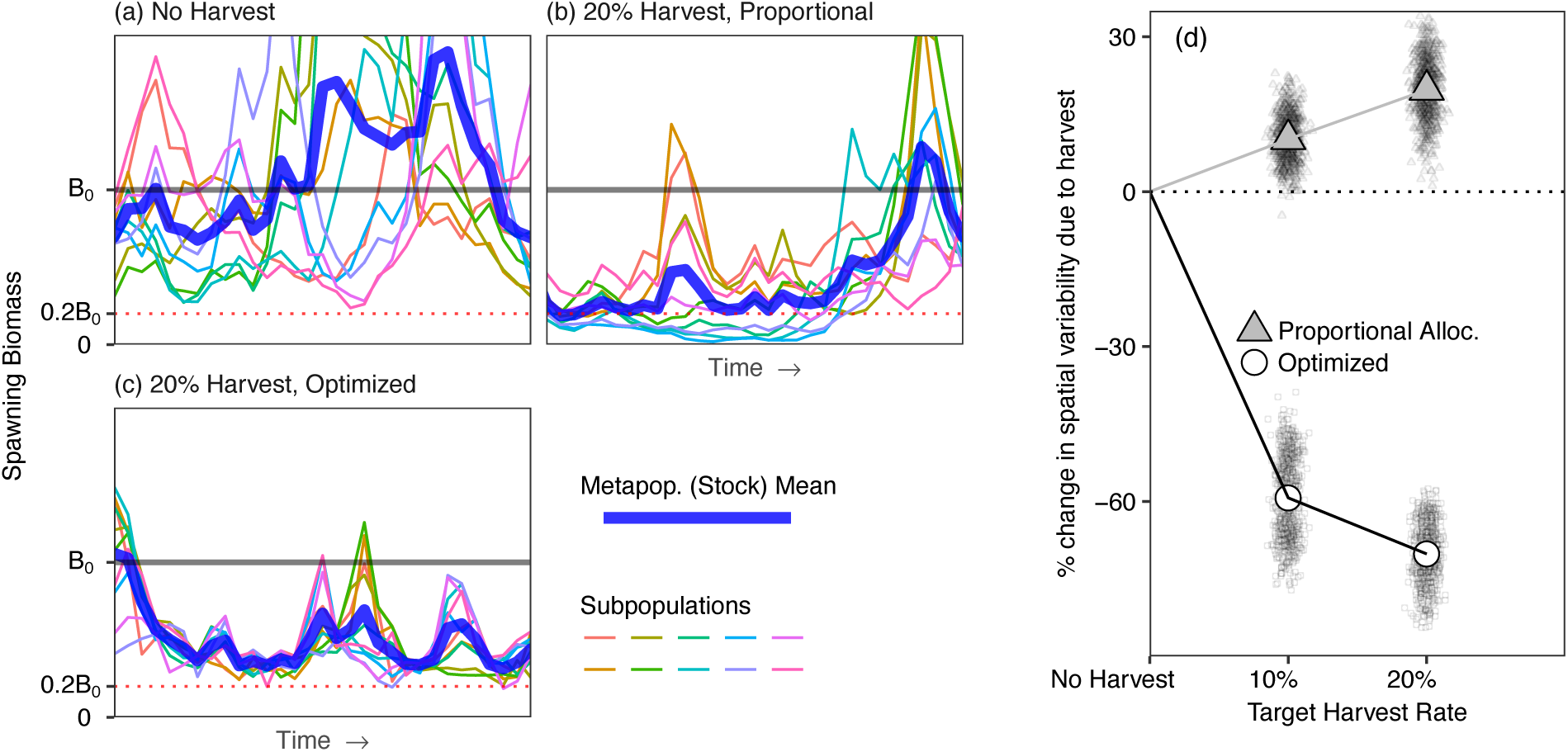
(a-c) Example simulations (a single simulation run) showing spatial variation in biomass measured at the subpopulation level (colored lines) or metapopulation mean (thick blue) for **(a)** an unexploited metapopulation, **(b)** a 20% target harvest allocated proportional to spawning biomass in space, or **(c)** a 20% target harvest allocated optimally in space according to the ideal free distribution (see Appendix S3: Fig. S4 for summaries across runs). B_0_ = deterministic biomass in the absence of fishing, dotted line = lower conservation threshold of 20% of B_0_. **(d)** change in spatial variation for different harvest rates and spatial harvest allocation across different spatial parameter scenarios for migration rates and spatial synchrony. Each point represents the spatial variance from 100 runs from a single 52-year simulation using a unique migration-synchrony parameter combination.

However, dynamically optimizing harvest rates in space to match local scale variability lowers the risk of subpopulation and aggregate metapopulation collapse (Fig. 3 d vs f and Fig. 3 h vs i) with no cost to aggregate catch (Appendix S3: Fig. S3). These results emerge from simulating a form of spatially optimized fishing effort that leaves underperforming locations unexploited and heavily targets overperforming local subpopulations, in accordance with predictions of ideal free distribution theory. Optimal spatial allocation of harvest rates reduces the effects of harvest intensity on spatial variability (Fig. 4d) which is strongly related to the bias in estimating risk among scales (Appendix S3: Fig. S1) and also reduces length of individual collapses (Appendix S3: Fig. S3). Thus, approaches that optimize spatial harvest to account for subpopulation dynamics can minimize the spatially destabilizing effect of harvest as well as disparity in exposure to risk of collapse among spatial scales. The benefits of spatially optimized harvest in terms of risk to subpopulations diminishes as migration and spatial synchrony in recruitment productivity decline to low levels (i.e. approach the lower left quadrant of panels in Fig. 3 d,f,h,j, where subpopulations are nearly autonomous with low connectivity).

## Discussion

Harvest strategies that appear appropriately prescribed at large spatial scales can, at local scales, lead to declines or even effective extirpation of local subpopulations. We call these small-scale declines “cryptic collapses”. Specifically, regional harvest strategies can create a “gilded trap” (Steneck et al. 2011) where, in this case, management focuses on the aspects of metapopulations that can benefit conservation and economics at the aggregate scale, but neglect social-ecological inequity in the exposure to risk at local scales. Our results show that spatial mismatch among scales is not merely an esoteric concern. Indeed, they occur in current management situations, and are supported by ecological theory we develop here. Our multiscale risk analysis highlights the impacts of scale mismatch on consumers and the potential value of optimized spatial management for sustainability and equity. Previous studies have also investigated spatial mismatches in fisheries contexts to understand consequences stemming from spatial mismatches among biological processes and available data, as well as the spatial implementation of fisheries for yield and measure of population status assessments (Cope and Punt 2011, McGilliard et al. 2011, Spies et al. 2015, McGilliard et al. 2017). Our work expands on previous investigations by considering how spatial dynamics of fish and fisheries affect resource sustainability at multiple spatial scales that are relevant to different species and fishing communities. Our models address this gap by building on existing research with increased biological realism to allow for 1) adult migration rates among subpopulations rather than only dispersal associated with recruitment (e.g. Cope and Punt 2011, McGilliard et al. 2011, Spies et al. 2015, McGilliard et al. 2017); 2) a range of complex spatio-temporal correlations in the stochastic populations dynamics (but see McGilliard et al. 2011 for an implementation of spatio-temporal variation in adult mortality); and 3) a novel suite of fisheries spatial harvest strategies.

### Insights into the benefits of population portfolios to manage risk for different groups

Our work adds critical resolution and understanding to the literature on portfolio effects that is focused on the benefits of spatial variability. Recent work viewing multiple populations as a “portfolio” of assets has shown benefits of maintaining a diversity of subpopulations with high asynchrony. These benefits include reducing local extinctions through so called ‘rescue effects’ (Hill et al. 2002, Secor et al. 2009, Fox et al. 2017), providing increased stability in the form of food security for people or animals (Nesbitt and Moore 2016) and minimizing economic risks over large scales by minimizing variance in harvestable abundance (Schindler et al. 2010). Yet these portfolio analyses typically focus on the attributes of the aggregate metapopulation, whereas the risk of localized subpopulation collapse (e.g. depletion below a threshold of ecological functionality or socioeconomic value) affects the interests of locally constrained fishers and spatially constrained organisms with small home ranges. Using theory and data we show that the same spatial variation that leads to resilience at the metapopulation scale, when left unaccounted for in management strategies, can also produce unforeseen negative consequences in the form of magnified spatial variation and local risk of collapse (see also Spies et al. 2015).

In the case of Pacific herring in the Central Coast of British Columbia, local reductions in some subpopulations occurred well before the entire stock showed substantial declines in the mid-2000s. These collapses had greater impact on spatially constrained groups; namely Indigenous communities for whom herring is a source of cultural and economic vitality (Brown and Brown 2009, Gavreau et al. 2017). In contrast, mobile fishing fleets and transient predators should be less vulnerable to local depletion events in the short term. Such context dependent effects of ignoring spatial variation are exemplified through considering the dramatic differences in the spatial scale at which predators and fishers operate. Indigenous fishers are spatially constrained by boat size, fuel costs, and political/cultural boundaries (Fig 2a, Table 4, Harris 2000, von der Porten et al. 2016). In contrast, the commercial fleet of seine and gillnet fishers are highly mobile and can pursue fish throughout the region. Similarly, non-human predators of herring and herring roe have a diversity of home-ranges and therefore interact with herring at multiple scales (Table 4). Many predators rely on herring when they move inshore around spawning season. Many crustaceans (Hines 1982, Stone and O’Clair 2001), echinoderms (Mattison et al. 1976, Cieciel et al. 2009), rockfishes and lingcod (Jorgensen et al. 2006, Mitamura et al. 2009, Tolimieri et al. 2009, Beaudreau and Essington 2011, Green and Starr 2011, Freiwald 2012), harbor seals (Peterson et al. 2012, Ward et al. 2012), some seabirds (Peery et al. 2009, Barbaree et al. 2015, Lorenz et al. 2017), and some flatfishes (Moser et al. 2013), exhibit restricted patterns of movement and are likely to exploit one to several major subpopulations, but generally not the entire spatial distribution of the metapopulation (stock). In contrast, humpback whales (Dalla Rosa et al. 2008, Kennedy et al. 2014), orcas (Hauser et al. 2007, Fearnbach et al. 2014), some seabirds (Pearce et al. 2005), sea lions (Merrick and Loughlin 1997, Fearnbach et al. 2014, Kuhn and Costa 2014), fur seals (Kuhn et al. 2014), halibut (Loher 2008, Seitz et al. 2011, Nielsen et al. 2014), and gadiforms (Wespestad et al. 1983, Hanselman et al. 2014, Rand et al. 2014) can, given ranges reported, access the geographic area covered by the stock (Table 4, see DataS1:Appendix S4). Thus, herring provide food resources to groups with varying movement constraints. As a result, herring collapses that range from small-scale subpopulations to metapopulation-wide phenomena may have differential impacts on the diverse predators that depend on this resource. These spatial scale dependencies affect which fishing communities or species bear the brunt of management risks and who reaps the benefits from the portfolio payoff of regional metapopulation stability. These outcomes highlight that ignoring space can exclude critical social, ecological, and economic responses central to the triple bottom line (Elkington 1994, Okamoto et al. 2019).

**Table 4:** Home-range categories of different Pacific herring predator groups. Details of sources and home-ranges are provided in the DataS1:Appendix S4.

| Home-Range Category | Home-Range Description | Herring Predator Groups |
| --- | --- | --- |
| Localized | single subpopulation<br>(*< 10 km radius*) | Indigenous fishers**, Seabirds*, Flatfishes*, Crustaceans*, Urchins, Cucumbers, Reef Fishes, Rockfishes |
| Centralized | multiple subpopulations<br>(*~10-40 km-radius*) | Indigenous fishers**, Seabirds*, Flatfishes*, Crustaceans*, Gadiforms*, Salmonids**, Halibut**, Harbor Seals |
| Transient | full stock or greater | Seine & Gillnet Fishers, Cetaceans, Sea Lions, Seabirds*, Gadiforms*, Salmonids**, Halibut** |

\*Varies by species, \*\* Varies by migration phase, location or individual

Pacific herring fisheries on Canada’s west coast provide an empirical case where the scale of regional stock assessments masked episodic local overexploitation and subpopulation collapses. Here, high local harvest rates appear to be commonplace even when local subpopulations are depleted presumably because of at least two key factors. First, schooling fish are easy to catch even at low abundances (Mackinson et al. 1997) and thus the quota is likely to be achieved even if the abundance of spawning fish in a particular location is small. Second, managers are challenged with fulfilling quotas with imperfect spatial information about spawning abundances. While high local harvest rates may have contributed to subsequent local collapses observed in this case study, quantifying other confounding demographics (i.e. spatial variation in stochastic adult mortality) are necessary to explicitly test this hypothesis. While recent local collapses may have occurred in the absence of fishing, fishing is very likely imposing higher subpopulation sensitivity to any environmental or biotic influence on recruitment or survival by reducing adult longevity thereby eroding an important buffer against recruitment volatility (Essington et al. 2015) that we show can contribute to spatial variation. Importantly, spatial variation may not only affect the localized groups. In the long term, assumptions of spatial homogeneity can generate biased estimates of total productivity (Takashina and Mougi 2015) that may lead to overly optimistic harvest strategies. Thus, high local harvest rates may produce sequential depletion over time that eventually erodes the principal of the stock with potential to generate collapse of the portfolio as a whole (Spies et al. 2015).

### Linking harvest dynamics to spatial variation in population dynamics

Importantly, spatial variation in population dynamics is not independent of harvest dynamics. Rather, we demonstrate numerically and analytically that spatial variation is likely to increase with harvest. This can occur in part because harvesting reduces spatial coupling. Higher mortality is known to reduce the abundance of older age classes (Barnett et al. 2017) and increase sensitivity to fluctuations in recruitment (Beddington and May 1977, Bjørnstad et al. 2004, Hsieh et al. 2006, Shelton and Mangel 2011, Okamoto et al. 2016). In spatially structured systems, such reduced longevity diminishes the synchronizing effect of migration and elevates sensitivity of subpopulations to localized environmental effects. However, if the distribution of harvest in space can be optimized to more adequately accommodate the spatial distribution of fish, we show how the overall portfolio can benefit at multiple scales. Here, demographic asynchrony (in this case asynchrony in recruitment productivity) can be maintained while minimizing subpopulation risk. Translated into practice, our simulation and analytical results highlight two non-mutually exclusive solutions that provide more equitable spreading of risk among scales. First, reductions in overall harvest rates can ameliorate biases in risk among scales. This occurs by reducing baseline levels of risk and reducing effects of harvest on spatial variability. Second, spatially optimizeing harvest allocations can minimize spatial variance and ease pressure on at-risk subpopulations, thereby reducing risk of local scale depletion without sacrificing catch. The first solution creates a trade-off between commercial yield and local risk; the second solution between the costs of management and fleet transportation, and local risk of subpopulation collapse. Specifically, achieving something similar to the latter (second solution) in a realistic setting is likely to require either some combination of greater investment in spatial monitoring, spatial stock assessments (Punt et al. 2018), and in season-management. Thus, moving in the direction of spatial optimization is likely to require substantial investment in costs and personnel for research, stock assessment, and management.

Addressing spatial inequity in risk exposure requires confronting these economic and logistical trade-offs. For species such as Pacific herring that have volatile spatiotemporal dynamics and complex migratory phenology (Benson et al. 2015), polycentric governance structures where governing authorities are nested at different spatial scales may help balance these trade-offs by addressing the problems of fit between ecosystems, social systems, and management agencies (Young 2002, Berkes 2006, Borgström et al. 2006, Folke et al. 2007, Biggs et al. 2012, von der Porten et al. 2016). Such systems can capitalize on scale-specific ecological knowledge (including local, traditional, and scientific knowledge), scientific capacity and socioeconomic experience to 1) guide decision analyses, 2) co-coordinate data collection and harvest allocation in space, and 3) test policies (e.g. in the Maine Lobster fishery (Acheson 2003)). Polycentric management schemes, however, are not a silver bullet. For systems like Pacific herring where placed-based Indigenous fishing communities often object to purely centralized scales of management for social, ecological, and economic reasons (Brown and Brown 2009, Thornton and Kitka 2015, von der Porten et al. 2016, Gavreau et al. 2017, von der Porten et al. 2019), opportunities to integrate knowledge and objectives into management strategies (and their evaluation) at smaller spatial scales is well placed in managing these fisheries, and may be critical to their perpetuity (Okamoto et al. 2019, Salomon et al. 2019).

The models used to generate inference in this study are simple in comparison to the nature of complex stochastic systems in space. The analytical model (Model 1) makes numerous simplifications (linearization, biological simplicity) in order to generate analytical and generalizeable solutions but may ignore more nuanced nonlinearities. The numerical model (Model 3) on the other hand is more detailed but outcomes are context dependent. Moreover, both models ignore many biological and management scenarios that are likely to further complicate spatial patterns (e.g. behaviorally or geographically complex migration (MacCall et al. 2018, Rogers et al. 2018), spatial and age specific mortality (McGilliard et al. 2011), spatially complex density dependence (McGilliard et al. 2017), Allee effects, cost, and data limitations for spatial management). We also ignore many alternative spatial allocation strategies that may be explicitly designed to maximize long-term yields or minimize spatiotemporal variability in part because these approaches would require a spatial assessment model, which is out of the scope of this study. However, the principles from our simulation and analytical models should generalize regardless of the degree of complexity in the system: a precautionary approach cognizant of resource users across multiple spatial scales may necessitate the incorporation of some degree of locally based management to minimize spatial discrepancies in risk exposure. Our results highlight the need to consider diverse scenarios and incorporate fundamental biological attributes that may impact resource dynamics and people at different scales. Overall, our models suggest a mixture of management scales may be key to selecting and coordinating harvest levels in space to navigate towards sustainable and equitable outcomes (Cope and Punt 2011, Biggs et al. 2012).

## Conclusion

Our analyses illustrate the importance of considering spatial dynamics for determining how to most effectively balance equity in management and conservation strategies aimed at achieving social, economic, and ecological outcomes (Halpern et al. 2013, Law et al. 2017). These issues are often ignored by centralized management and conservation initiatives focused on larger spatial scales. For over half a century, fisheries scientists have debated how best to exploit and conserve “mixed stocks” that have separate dynamics but are inseparable in space, with the aim of balancing conservation of weak stocks and maximizing total yields (Ricker 1958). Our work inverts this focus to metapopulations where subpopulations have inseparable dynamics but are separated in space. We show how, in these settings, exploitation can magnify local scale fluctuations and spatial variability. As a result, aggregate metrics poorly represent local scale dynamics and the risk of collapse at the local scale increases at a greater rate than the aggregate large scale. This phenomenon creates cryptic collapses with ensuing discrepancies in risk exposure. Fortunately, the magnitude of these discrepancies can potentially be controlled by lowering harvest rates or seeking harvest dynamics that are spatially optimized. Overall, these conclusions are relevant not only to Pacific herring fisheries, but also to the great number of exploited natural resources that exhibit spatial structure and are valuable to species and people that operate on multiple scales.

## Supporting information

Metadata_Appendix S4

## Acknowledgements

DKO was supported by a Strategic Partnership Grant from the National Sciences and Engineering Research Council (NSERC) of Canada to AKS and MHL and a first-year assistant professor (FYAP) award to DKO from the FSU Council on Research and Creativity. We thank the Gordon and Betty Moore Foundation for their support of AS, JSF, PSL, the Ocean Tipping Points project and the David and Lucille Packard Foundation for their support of PSL, the Tula Foundation for their support of MHL, and the Ocean Modeling Forum. We thank J. Cleary, S. Harper, M. Reid, K. Gladstone, B. Gladstone, K. Brown, and D. Neasloss for discussions that initiated and refined the research, A. Frid and A. Rassweiler for input on the manuscript, and S. Cox, L. Dee, L. Hauser, B. Hunt, S. Miller, I. McKechnie, S. Pau, T. Pitcher, E. Petrou, T. Francis, A. Punt and W. Smith for discussions that improved the research. We also thank the Heiltsuk and Haida First Nations for their partnership in the NSERC Strategic Partnership Grant and the Department of Fisheries and Oceans for providing herring spatial data.

