## Appendix S1 for "Spatial variation in exploited metapopulations obscures risk of collapse"

### Appendix S1: Derivation of the VAR(1) model & properties
### "Spatial variation in exploited metapopulations obscures risk of collapse"
### *Ecological Applications*

Daniel K Okamoto[a,b,c], Margot Hessing-Lewis[b], Jameal F Samhouri[d], Andrew O Shelton[d], Adrian Stier[e], Philip S Levin[f,g], and Anne K Salomon[b,c]

[a]Department of Biological Science, Florida State University, Tallahassee, Florida 32303, USA
[b]Hakai Institute, PO Box 309, Heriot Bay, BC V0P 1H0, Canada
[c]School of Resource and Environmental Management, Simon Fraser University, 8888 University Drive, Burnaby, B.C. V5A 1S6, Canada
[d]Conservation Biology Division, Northwest Fisheries Science Center, National Marine Fisheries Service, National Oceanic and Atmospheric Administration, 2725 Montlake Blvd. East,Seattle, WA 98112, USA
[e]Department of Ecology, Evolution, and Marine Biology, University of California, Santa Barbara, CA 93106, USA
[f]The Nature Conservancy, 74 Wall St, Seattle, WA98121, USA
[g]School of Environment and Forestry Sciences, University of Washington, Seattle, Washington, USA

**Corresponding Author**
Daniel K Okamoto
Department of Biological Science
The Florida State University
Tallahassee, Florida 32303


**Author contributions**

DKO designed, built and conducted analyses. AS conducted the home-range literature review. DKO and AKS wrote the initial manuscript draft. DKO, MHL, JFS, AOS, AS, PSL, and AKS initiated the research focus, refined analyses and contributed to revisions.

### Appendix S1: Derivation of the VAR(1) model & properties

*Deriving the Jacobian*

Each row of the Jacobian matrix is a vector of partial derivatives of each age class within a given subpopulation with respect to each other age class within and among subpopulations. Thus, the $k$th element of the $j$th row in $J_i$ is the partial derivative of age class $j$ with respect to age class $k$ within each subpopulation (for $J_i = J_1$) or across subpopulations (for $J_i = J_2$). Below we derive the estimates for within subpopulation entries ($J_i = J_1$). The same calculations apply for among subpopulation entries by replacing $\delta$ with $(1-\delta)$ and vice versa.

Zygotes are only produced by age 1+ and age classes are produced by the prior age class in the year prior at the two subpopulations. On the log-scale, this translates to:

$$x_{a=0,i} = \ln(e^{-Z}[(1-\delta)\sum_{a=1}^{n} e^{x_{a,i}} + (\delta)\sum_{a=1}^{n} e^{x_{a,j}}]) \tag{S1}$$

The partial derivative of log-zygote producers with respect to each age class $a$ (not including $x_{0,i}$) is produced by the chain rule:

$$\frac{\partial x_{a=0,i}}{\partial x_{a,i}} = e^{-Z}(1-\delta)e^{x_{a,i}}/(e^{-Z}[(1-\delta)\sum_{a=1}^{n} e^{x_{a,i}} + (\delta)\sum_{a=1}^{n} e^{x_{a,j}}]) \tag{S2a}$$

$$= (1-\delta)e^{x_{a,i}}/[(1-\delta)\sum_{a=1}^{n} e^{x_{a,i}} + (\delta)\sum_{a=1}^{n} e^{x_{a,j}}] \tag{S2b}$$

Setting the equilibrium values for each age class to:

$$e^{x_a^*} = e^{x_{a-1}^*}e^{-Z} = e^{x_{a-2}^*}(e^{-Z})^2 = ... = e^{x_{a=1}^*}(e^{-Z})^{a-1} \tag{S3}$$

yields the entries for the first row of the Jacobian that is independent of the state variables:

$$\frac{\partial x_{a=0,i}}{\partial x_{a,i}} = (1-\delta)e^{x_{a=1}^*}(e^{-Z})^{a-1}/(e^{x_{a=1}^*} + e^{x_{a=1}^*}\sum_{a=1}^{n}(e^{-Z})^{a-1}) \tag{S4a}$$

$$= (1-\delta)(e^{-Z})^{a-1}/(1 + \sum_{a=1}^{n}(e^{-Z})^{a-1}) \tag{S4b}$$

The partial derivative of age 1 individuals with respect to zygotes is determined by:

$$x_{a=1,i} = \ln\alpha + (1-\beta)x_{a=0,i}; \qquad \frac{\partial x_{a=1,i}}{\partial x_{a=0,i}} = (1-\beta) \tag{S5}$$

The partial derivative of age a individuals with respect to age a-1:

$$x_{a,i} = \ln(e^{-Z}(1-\delta)e^{x_{a-1,i}} + e^{-Z}(\delta)e^{x_{a-1,j}})\delta x_{a-1,i} \tag{S6}$$

$$\begin{aligned}\frac{\partial x_{a,i}}{\partial x_{a-1,i}} &= e^{-Z}(1-\delta)e^{x_{a-1,i}}/(e^{-Z}(1-\delta)e^{x_{a-1,i}} + e^{-Z}(\delta)e^{x_{a-1,j}}) && \text{(S7a)}\\ &= e^{-Z}(1-\delta)e^{x^*_{a-1}}/(e^{-Z}e^{x^*_{a-1}}) && \text{(S7b)}\\ &= (1-\delta) && \text{(S7c)}\end{aligned}$$

Finally, partial derivative of the plus group individuals with respect to age $a = n-1$ or $a = n$ is again given by the chain rule:

$$x_{n,i} = \ln(e^{-Z}(1-\delta)(e^{x_{a-1,i}} + e^{x_{a,i}}) + e^{-Z}(\delta)(e^{x_{a-1,j}} + e^{x_{a-1,j}}))\delta x_{a-1,i} \tag{S8}$$

$$\begin{aligned}\frac{\partial x_{n,i}}{\partial x_{a,i}} &= e^{-Z}(1-\delta)(e^{x_{a,i}})/(e^{-Z}(\delta-1)(e^{x_{n-1,i}} + e^{x_{n,i}}) + (\delta)(e^{x_{n-1,j}} + e^{x_{n,j}})) && \text{(S9a)}\\ &= (1-\delta)(e^{x_{a,i}})/((\delta-1)(e^{x_{n-1,i}} + e^{x_{n,i}}) + (\delta)(e^{x_{n-1,j}} + e^{x_{n,j}})) && \text{(S9b)}\end{aligned}$$

$$\frac{\partial x_{n,i}}{\partial x_{n-1,i}} = (1-\delta)/(1+e^{-Z}) \tag{S10}$$

$$\frac{\partial x_{n,i}}{\partial x_{n,j}} = (1-\delta)e^{-Z}/(1+e^{-Z}) \tag{S11}$$

*Deriving the Statistical Properties*

The long-term temporal variability of the individual populations and the metapopulation is determined by the long-run covariance matrix of a VAR(1) (Lütkepohl 2005):

$$\mathrm{vec}(\mathbf{\Sigma}_x) = [\boldsymbol{I}_{p^2} - \boldsymbol{J} \otimes \boldsymbol{J}]^{-1}\,\mathrm{vec}(\mathbf{\Sigma}_\zeta) \tag{S12}$$

where $p$ is the total number of variables (i.e. zygotes and age classes in both subpopulations), $\boldsymbol{I}_{p^2}$ is a $p^2 \times p^2$ identity matrix, $\otimes$ is the Kronecker product, and vec is the vector operator (i.e. turning a matrix into a vector for operational sake). The population and metapopulation coefficient of variation (on the natural scale), as well as the among population spatial variation and spatial correlations the can be derived from $\mathbf{\Sigma}_x$ using the definition of the mean, variance and covariance of a multivariate lognormal given the mean and covariance matrix of the log-scale system. Derivations are shown below.

If we assume the system conforms to a multivariate lognormal, then the temporal variance of spawners at site $i$ is:

$$\mathrm{Var}(e^{[x_{i,a=0}]}) = e^{[2\bar{x}_{i,a=0}+\sigma^2_{i,i}]}e^{[\sigma^2_{i,i}-1]} = e^{[\sigma^2_{i,i}]}e^{[\sigma^2_{i,i}-1]} \tag{S13}$$

where $\sigma^2_{i,i}$ is the diagonal element of the log-scale variance covariance matrix $\mathbf{\Sigma}_x$ associated with spawners at site $i$ ($x_{i,a=0}$), and $\bar{x}_{i,a=0}$ is the mean of $x_{i,a=0}$ which is set to zero. From the lognormal, the mean of $e^{[x_{i,a=0}]}$ is $e^{[0.5\sigma^2_{i,i}]}$.

The resulting coefficient of variation of each spawning population is:

$$\mathrm{CV}(e^{[x_{i,a=0}]}) = \frac{\sqrt{e^{\left[\sigma^2_{i,i}\right]}e^{\left[\sigma^2_{i,i}-1\right]}}}{e^{\left[0.5\sigma^2_{i,i}\right]}} = \sqrt{e^{\left[\sigma^2_{i,i}-1\right]}} \tag{S14}$$

The temporal covariance of scaled spawners among sites is:

$$\mathrm{Cov}(e^{[x_{i,a=0}]}, e^{[x_{j,a=0}]}) = e^{[0.5\sigma^2_{i,i}+0.5\sigma^2_{j,j}]}e^{[\sigma^2_{i,j}-1]} \tag{S15}$$

where $\sigma^2_{i,j}$ is the off-diagonal (covariance) element of $\mathbf{\Sigma}_x$.

Considering a metapopulation with two component populations, we set the log-population variances equal (i.e. $\sigma_{i,i} = \sigma_{j,j} = \sigma$), the resulting variance of the spatial mean on the natural scale is:

$$\mathrm{Var}\left[\frac{e^{[x_{i,a=0}]} + e^{[x_{j,a=0}]}}{2}\right] = \mathrm{Var}\left(\frac{e^{[x_{i,a=0}]}}{2}\right) + \mathrm{Var}\left(\frac{e^{[x_{j,a=0}]}}{2}\right) + 2\mathrm{Cov}\left(\frac{e^{[x_{i,a=0}]}}{2}, \frac{e^{[x_{j,a=0}]}}{2}\right) \tag{S16}$$

$$= 0.5e^{[\sigma^2]}e^{[\sigma^2-1]} + 0.5e^{[\sigma^2]}e^{[\sigma^2_{i,j}-1]} \tag{S17}$$

Thus, the temporal coefficient of variation of the spatial mean is:

$$\mathrm{CV}\left[\frac{e^{[x_{i,a=0}]} + e^{[x_{j,a=0}]}}{2}\right] = \frac{\sqrt{0.5e^{[\sigma^2]}e^{[\sigma^2-1]} + 0.5e^{[\sigma^2]}e^{[\sigma^2_{i,j}-1]}}}{e^{[0.5\sigma^2]}} = \sqrt{0.5e^{[\sigma^2-1]} + 0.5e^{[\sigma^2_{i,j}-1]}} \tag{S18}$$

In this case, the residual spatial variance (subtracting S.17 from S.13 according to S.32 below) is:

$$\text{residual spatial variance} = 0.5e^{[\sigma^2]}e^{[\sigma^2-1]} - 0.5e^{[\sigma^2]}e^{[\sigma^2_{i,j}-1]} \tag{S19}$$

Which when standardized by the square of the population mean ($e^{[0.5\sigma^2]}$) yields:

$$\text{standardized residual spatial variance} = 0.5e^{[\sigma^2-1]} - 0.5e^{[\sigma^2_{i,j}-1]} \tag{S20}$$

In this case this is the same as the variance of the difference between either population and the population mean trend (because both populations are identical).

$$\mathrm{Var}\left[e^{[x_{i,a=0}]} - \left(\frac{e^{[x_{i,a=0}]} + e^{[x_{j,a=0}]}}{2}\right)\right] = \mathrm{Var}\left[\frac{e^{[x_{i,a=0}]}}{2} - \frac{e^{[x_{j,a=0}]}}{2}\right] \tag{S21}$$

$$= \mathrm{Var}\left[\frac{e^{[x_{i,a=0}]}}{2}\right] + \mathrm{Var}\left[\frac{e^{[x_{j,a=0}]}}{2}\right] - 2\mathrm{Cov}\left(\frac{e^{[x_{i,a=0}]}}{2}, \frac{e^{[x_{j,a=0}]}}{2}\right) \tag{S22}$$

$$= 0.5e^{[\sigma^2]}e^{[\sigma^2-1]} - 0.5e^{[\sigma^2]}e^{[\sigma^2_{i,j}-1]} \tag{S23}$$

As a result, we can generate the residual spatial variation for the metapopulation and how it is influenced by mortality and environmental correlations.

*Autocorrelations*

The matrix of first-order autocorrelations that yields within population correlation and spatial coupling is given by:

$$\mathbf{R} = (\mathrm{diag}(\mathbf{\Sigma}_x)^{\circ\frac{1}{2}})^{-1}\mathbf{\Sigma}_x(\mathrm{diag}(\mathbf{\Sigma}_x)^{\circ\frac{1}{2}})^{-1} \tag{S24}$$

*Impulse response of the VAR(1) model*

The impulse response (IR) in a VAR(1) model are given by the moving average (MA) representation of the VAR model at a given lag. Specifically, the matrix of MA coefficients for the VAR(1) at lag $t$ is produced raising the matrix of VAR(1) coefficients to the $t^{th}$ power. The first order impulse response is thus simply the matrix of VAR(1) coefficients, which in our case is conveniently given by the jacobian matrix $\mathbf{J}$. We use a first-order impulse response because the greatest IR of adults is at lag 1 (when a recruitment pulse first enters into the adult stage). In our case, the variables subject to stochasticity in recruitment at a single lag are the new adults at location $i$, which in this case is the 2nd column of the 1st row in $J_1$ defined in Eq. 5. Because this represents the log-scale additive response, the IR on the natural scale is a multiplicative effect.
