## Appendix S2 for "Spatial variation in exploited metapopulations obscures risk of collapse"

### Appendix S2: Substock trends in Central Coast herring
### "Spatial variation in exploited metapopulations obscures risk of collapse"
*Ecological Applications*

Daniel K Okamoto[a,b,c], Margot Hessing-Lewis[b], Jameal F Samhouri[d], Andrew O Shelton[d], Adrian Stier[e], Philip S Levin[f,g], and Anne K Salomon[b,c]

[a]Department of Biological Science, Florida State University, Tallahassee, Florida 32303, USA
[b]Hakai Institute, PO Box 309, Heriot Bay, BC V0P 1H0, Canada
[c]School of Resource and Environmental Management, Simon Fraser University, 8888 University Drive, Burnaby, B.C. V5A 1S6, Canada
[d]Conservation Biology Division, Northwest Fisheries Science Center, National Marine Fisheries Service, National Oceanic and Atmospheric Administration, 2725 Montlake Blvd. East,Seattle, WA 98112, USA
[e]Department of Ecology, Evolution, and Marine Biology, University of California, Santa Barbara, CA 93106, USA
[f]The Nature Conservancy, 74 Wall St, Seattle, WA98121, USA
[g]School of Environment and Forestry Sciences, University of Washington, Seattle, Washington, USA

**Corresponding Author**
Daniel K Okamoto
Department of Biological Science
The Florida State University
Tallahassee, Florida 32303


**Author contributions**
DKO designed, built and conducted analyses. AS conducted the home-range literature review. DKO and AKS wrote the initial manuscript draft. DKO, MHL, JFS, AOS, AS, PSL, and AKS initiated the research focus, refined analyses and contributed to revisions.

#### Appendix S2: Substock trends in Central Coast herring

*Residual spatial variability*

The residual spatial variation (RSV, also called $\beta$ variation or the spatial variation due to asynchrony) is the difference between the variance of the mean metapopulation trend and the expected metapopulation mean variance if populations are perfectly correlated. In other words, residual spatial variability is the additional temporal variability at the subpopulation level left unexplained by the temporal variability of the metapopulation mean.

The variability of a metapopulation mean given individual variability and covariance of each of $N$ individual population time series ($X_i$) is given by:

$$\mathrm{Var}\left(\frac{1}{N}\sum_{i=1}^{N} X_i\right) = \frac{1}{N^2}\left[\sum_{i=1}^{N}\mathrm{Var}\left(X_i\right) + \sum_{i=1}^{N}\sum_{j\neq 1}^{N-1}\rho_{i,j}\sqrt{\mathrm{Var}\left(X_i\right)\mathrm{Var}\left(X_j\right)}\right] \tag{S1}$$

$$= \frac{1}{N^2}\left[\sum_{i=1}^{N}\mathrm{Var}\left(X_i\right) + \sum_{i=1}^{N}\sum_{j\neq 1}^{N-1}\mathrm{Cov}\left(X_i, X_j\right)\right] \tag{S2}$$

$$\tag{S3}$$

In contrast, the maximum potential variance of a metapopulation mean given individual variance of each of $N$ populations is:

$$\mathrm{Var}_{\max}\left(\bar{X}\right) = \frac{1}{N^2}\left[\sum_{i=1}^{N}\mathrm{Var}\left(X_i\right) + \sum_{i=1}^{N}\sum_{j\neq 1}^{N-1}\sqrt{\mathrm{Var}\left(X_i\right)\mathrm{Var}\left(X_j\right)}\right] = \left(\sum_{i=1}^{N}\frac{\sqrt{\mathrm{Var}\left(X_i\right)}}{N}\right)^2 \tag{S4}$$

Hence, the residual spatial variance (RSV) is:

$$\mathrm{RSV}\left(\bar{X}\right) = \mathrm{Var}_{\max}\left(\bar{X}\right) - \mathrm{Var}\left(\bar{X}\right) \tag{S5}$$

$$= \frac{1}{N^2}\left[\sum_{i=1}^{N}\sum_{j\neq 1}^{N-1}\sqrt{\mathrm{Var}\left(X_i\right)\mathrm{Var}\left(X_j\right)} - \sum_{i=1}^{N}\sum_{j\neq 1}^{N-1}\mathrm{Cov}\left(X_i, X_j\right)\right] \tag{S6}$$

$$= \frac{1}{N^2}\left[\sum_{i=1}^{N}\sum_{j\neq 1}^{N-1}\left(1-\rho_{i,j}\right)\sqrt{\mathrm{Var}\left(X_i\right)\mathrm{Var}\left(X_j\right)}\right] \tag{S7}$$

$$= \left(\sum_{i=1}^{N}\frac{\sqrt{\mathrm{Var}\left(X_i\right)}}{N}\right)^2 - \mathrm{Var}\left(\bar{X}\right) \tag{S8}$$

Similarly, population synchrony as defined by X is given by:

$$Synchrony = \frac{\mathrm{Var}\left(\bar{X}\right)}{\mathrm{Var}\left(\bar{X}\right) + \mathrm{RSV}\left(\bar{X}\right)} \tag{S9}$$

*Model estimation and validation*

We estimated core parameters in the following manner :

1. Process error is multivariate lognormally distributed about the vector of pre-harvest biomass expectations with an estimated variance-covariance matrix. We estimated the spatial correlation in process error via the Cholesky decomposition of an unconstrained correlation matrix with an LKJ prior (Lewandowski et al. 2009). We estimate log-biomass knowing that a) pre-harvest biomass must be greater than or equal to the harvest, and b) process error affects dynamics before harvest. Thus, the conditional prior distribution for biomass is placed on logarithm of the sum of post-harvest biomass and harvest (Table 1, Eq. 11), conditional on the deterministic model expectations (given estimated spawning biomass from the year prior) and process error covariance matrix. Because of this change of variables (i.e. the prior is placed on a transformation) the observation likelihood is necessarily adjusted by the log-Jacobian of the inverse transform (Table 1, Eqs 11 & 12; Carpenter et al. 2016). Because the transform is in the logarithmic form, we use the chain rule to differentiate the logarithm of the sum, and then to differentiate the sum.

2. The deterministic parameters (biomass specific analogue to survival, density dependence, and density independent productivity) are given a hierarchical structure; the vector of mean parameters are assumed to be correlated with a multivariate normal prior, and the site-specific parameters are also given a multivariate normal prior with the aforementioned mean vector, independent half-cauchy standard deviations and an LKJ prior for the Cholesky decomposition of the correlation matrix (Table 1).

3. We assumed egg surveys were never fully observed but provide a directionally biased index, consistent with the current stock assessment assumptions (Table 1); we impose on the log-bias (catchability) parameter a normal prior as in Martell et al. (2012).

4. We use a log-likelihood function that is the sum of the log-scale lognormal density given model expectations and the Jacobian adjustment from (1) above (Table 1, Eq. 11). All equations and priors used are shown in Table 1 .

52 *Full time series results*

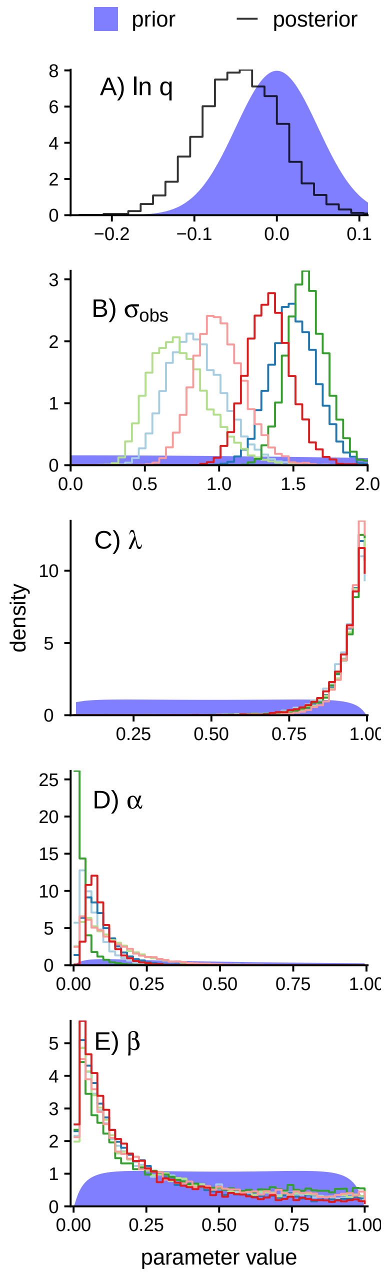


**Figure S1:** Posterior histograms (lines) and prior densities (blue bands) for key parameters in the model. Priors were chosen to be vague, with the exception of the survey bias (ln q, also known as a "catchability" for the survey) which was derived from the assumptions in the current assessment. Parameters include ln q (survey bias), $\sigma_{obs}$ (the site specific lognormal observation error), $\lambda$ (the aggregate parameter for survival and somatic growth), $\alpha$ (the Gompertz productivity parameter) and $\beta$ (the Gompertz compensation parameter). Priors are defined in Table 1.

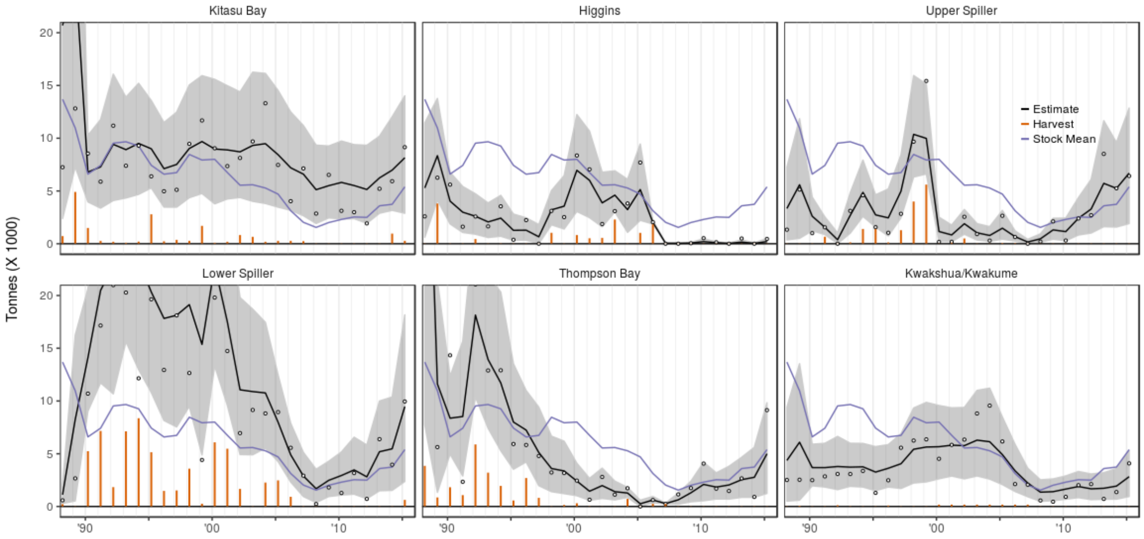


**Figure S2:** Trends in pre-spawning biomass (spawn plus catch) and associated catch for the six main spawning locations of Pacific herring the Central Coast of British Columbia. Open points represent the observed pre-spawn biomass, black lines represent posterior estimate of biomass from the model, blue lines represent the overall stock mean, and grey bands are the 10th and 90th% posterior quantiles of estimated biomass.

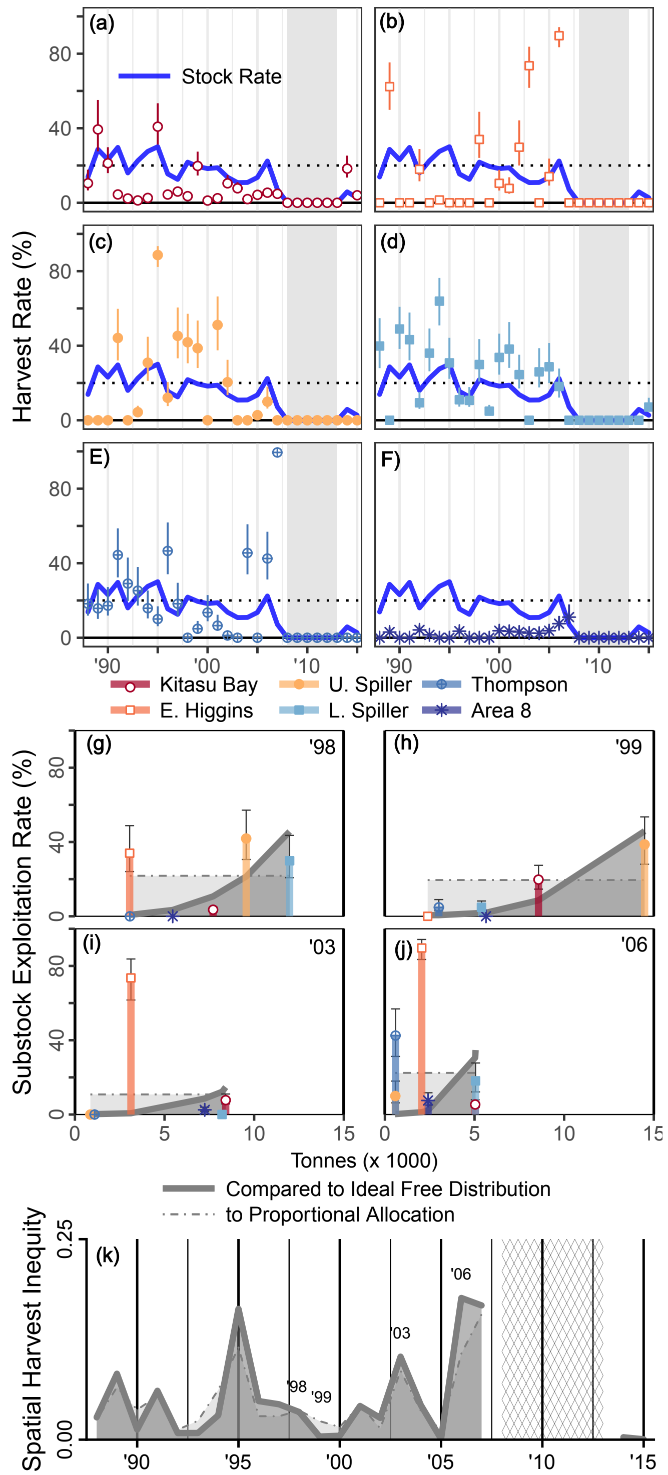


**Figure S3:** (a-f) Trends in annual harvest rate for the 6 subpopulations (substocks - points and 95% credibility intervals) and the mean for the metapopulation (stock - thick blue line). The dotted line represents the target harvest rate of 20%. (g-j) Harvest rate and 95% credible intervals for each section in four select years compared to the optimized strategy via the ideal free distribution of harvest (IFD, grey band) or stock allocation with uniform proportional harvest (broken grey line). Points above both the IFD and the equal stock allocation may represent a suboptimal harvest strategy. In contrast, points along the IFD line, such as that in 1999, represent a strategy may reduce local risks. (k) Spatial harvest inequity (measured as the absolute mean relative error from perfect equity). Note that in 1999 equity differs depending on which benchmark is used (IFD or uniform proportional harvest).

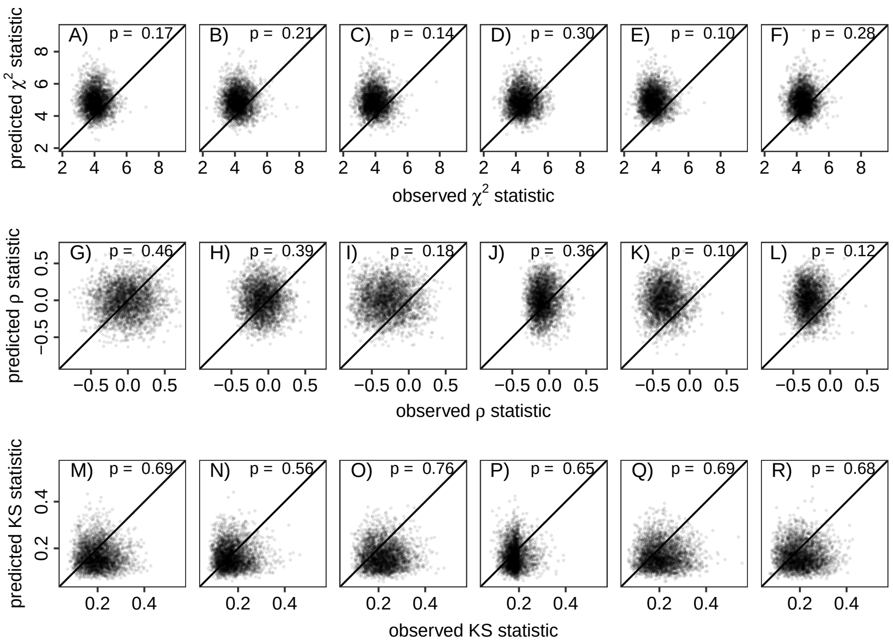


**Figure S4:** Posterior predictive checks for the hierarchical state-space model for three different statistics (rows) by site (columns). Observed statistics that exhibit dramatic departures from the predicted (i.e. the point cloud is significantly above or below the 1:1 line) suggest poor fit or model adequacy. P-values less than 0.05 or greater than 0.95 represent the a 95% probability of more or less extreme values than predicted. (A-F) Observed versus predicted $\chi^2$ goodness of fit test statistic. (G-J) Observed versus predicted Spearman's $\rho$ test statistic measuring correlation between model residuals and predicted biomass. (M-R) Observed versus predicted Kolmogorov-Smirnov test comparing model residuals to predicted under log-normal observation likelihood.
