## Appendix S3 for "Spatial variation in exploited metapopulations obscures risk of collapse"

Appendix S3: Simulation model details and full results
"Spatial variation in exploited metapopulations obscures risk of collapse"
*Ecological Applications*

Daniel K Okamoto[a,b,c], Margot Hessing-Lewis[b], Jameal F Samhouri[d], Andrew O Shelton[d], Adrian Stier[e], Philip S Levin[f,g], and Anne K Salomon[b,c]

[a]Department of Biological Science, Florida State University, Tallahassee, Florida 32303, USA
[b]Hakai Institute, PO Box 309, Heriot Bay, BC V0P 1H0, Canada
[c]School of Resource and Environmental Management, Simon Fraser University, 8888 University Drive, Burnaby, B.C. V5A 1S6, Canada
[d]Conservation Biology Division, Northwest Fisheries Science Center, National Marine Fisheries Service, National Oceanic and Atmospheric Administration, 2725 Montlake Blvd. East,Seattle, WA 98112, USA
[e]Department of Ecology, Evolution, and Marine Biology, University of California, Santa Barbara, CA 93106, USA
[f]The Nature Conservancy, 74 Wall St, Seattle, WA98121, USA
[g]School of Environment and Forestry Sciences, University of Washington, Seattle, Washington, USA

**Corresponding Author**
Daniel K Okamoto
Department of Biological Science
The Florida State University
Tallahassee, Florida 32303


**Author contributions**
DKO designed, built and conducted analyses. AS conducted the home-range literature review. DKO and AKS wrote the initial manuscript draft. DKO, MHL, JFS, AOS, AS, PSL, and AKS initiated the research focus, refined analyses and contributed to revisions.

### Appendix S3: Simulation model details and full results

*Spatial allocation of harvest*

The fishing fleet either distributes itself in space or a manager forces spatial allocation. We use three scenarios for spatial allocation: 1) diffuse effort in space (i.e harvest is proportional to abundance), 2) ideal distribution of effort in space, and 3) random spatial allocation.

- *Proportional allocation of harvest*

  For a proportional allocation of harvest, the calculation of biomass caught in each location ($\mathrm{H}_{l,t}$) is simply:

$$H_{\mathrm{adults}_{l,t}} = \frac{\sum_{a=2}^{10} \mathrm{w}_a \mathrm{m}_a n_{a,l,t}}{\sum_{l=1}^{L} \sum_{a=2}^{10} \mathrm{w}_a \mathrm{m}_a n_{a,l,t}} \times \mathrm{Quota}_t \tag{S1}$$

- *Optimized Allocation*

  Optimized spatial allocation is generated using the ideal free distribution of effort in space. From an efficiency standpoint, the fishing fleet would allocate themselves to optimize efficiency of catch (i.e. minimal effort for the return). We can numerically solve for the spatial harvest strategy that optimizes efficiency (assuming no movement costs and perfect information) using a system of ordinary differential equations (ODEs). First, we 1) simplify the fishery to a very short time period, 2) assume vessels can move instantaneously among locations with no cost, 3) assume vessels have perfect information and 4) assume fishing effort (i.e. sets) can be broken into infinitesimally small units. Consider $F_{l,\tau}$ the rate of fishing effort (i.e. number of sets per unit time) allocated at time $\tau$ in location $l$, $q$ is the catchability of fish (i.e. ratio of proportional capture rate to effort), $B_\tau$ is the biomass available for capture at time $\tau$ $D_e$ is the rate at which vessels respond to spatial disparities in efficiency, where efficiency = effort x catchability x biomass. To maximize efficiency, vessels sense disparities in efficiency and move instantaneously to neighboring locations with higher efficiency (i.e. catch per unit time per unit effort).

$$\frac{dB_l}{d\tau} = -F_l(\tau) q B_l(\tau) \tag{S2a}$$

$$\frac{dF_l}{d\tau} = \sum_{j=1}^{L} \left[F_l(\tau) q B_l(\tau) - F_j(\tau) q B_j(\tau)\right] \tag{S2b}$$

$$\frac{dH_l}{d\tau} = F_l(\tau) q B_l(\tau) \tag{S2c}$$

$$\mathrm{H}(\tau_Q, l) = \int_0^{\tau_Q} F_l(\tau) q B_l(\tau) d\tau \tag{S3a}$$

$$\mathrm{StockH}(\tau_Q) = \sum_{l=1}^{L} \mathrm{H}(\tau_Q, l) \tag{S3b}$$

We numerically evaluate this system of ODEs until the quota is achieved to produce the final, optimal distribution of catch in space.

- *Alternative allocation strategies: Random spatial allocation distribution*

  We consider a less ideal scenario where vessels harvest at random from patches. This may occur when fleets can still effectively harvest from subpopulations at low biomass because fish school. For random distribution of the harvest, the proportion of the quota removed from each location within each year is a random draw from a Dirichlet distribution, with mean equal to the actual proportional biomass distribution. For this scenario, we limit harvest to a maximum of 90% at any given site. The fleet randomly chooses a site, prosecutes a fishery, and moves on until the quota is achieved. Functionally, the fleet effort is distributed randomly in space, with mean allocation probabilities given by the distribution of biomass in space. Potential, proportional harvest rates are randomly assigned to each spawning area (though cannot exceed the maximum) and prosecuted in order from highest to lowest until the quota is achieved. The initial random proportional harvest rates are set by a Dirichlet distribution where means are proportional to the distribution of stock biomass in space and a count parameter ($K$) that sets the degree of randomness (set as $K = 5$) to achieve a spatial harvest inequity of approximately 0.20. Lower count parameters defy the proportionality more (i.e. harvests are unlikley to reflect biomass distributions in space) while very high count parameters approximate harvests that are porportional to biomass distributions in space.

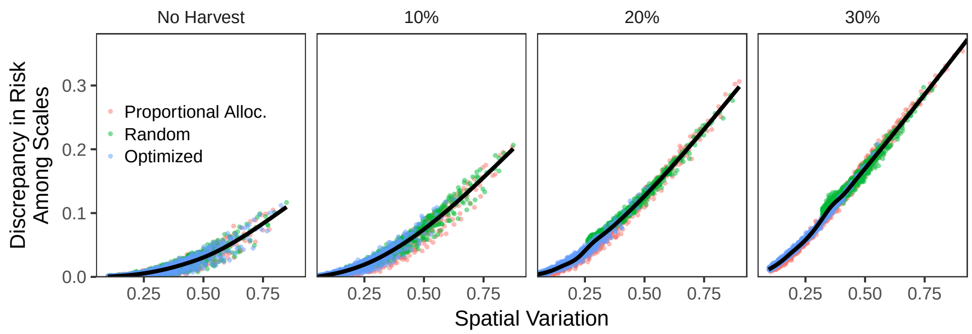


**Figure S1:** Discrepancy in risk among spatial scales (mean risk at subpopulation scale minus mean risk at metapopulation scale) versus spatial variance by harvest rate and spatial harvest allocation strategy

.

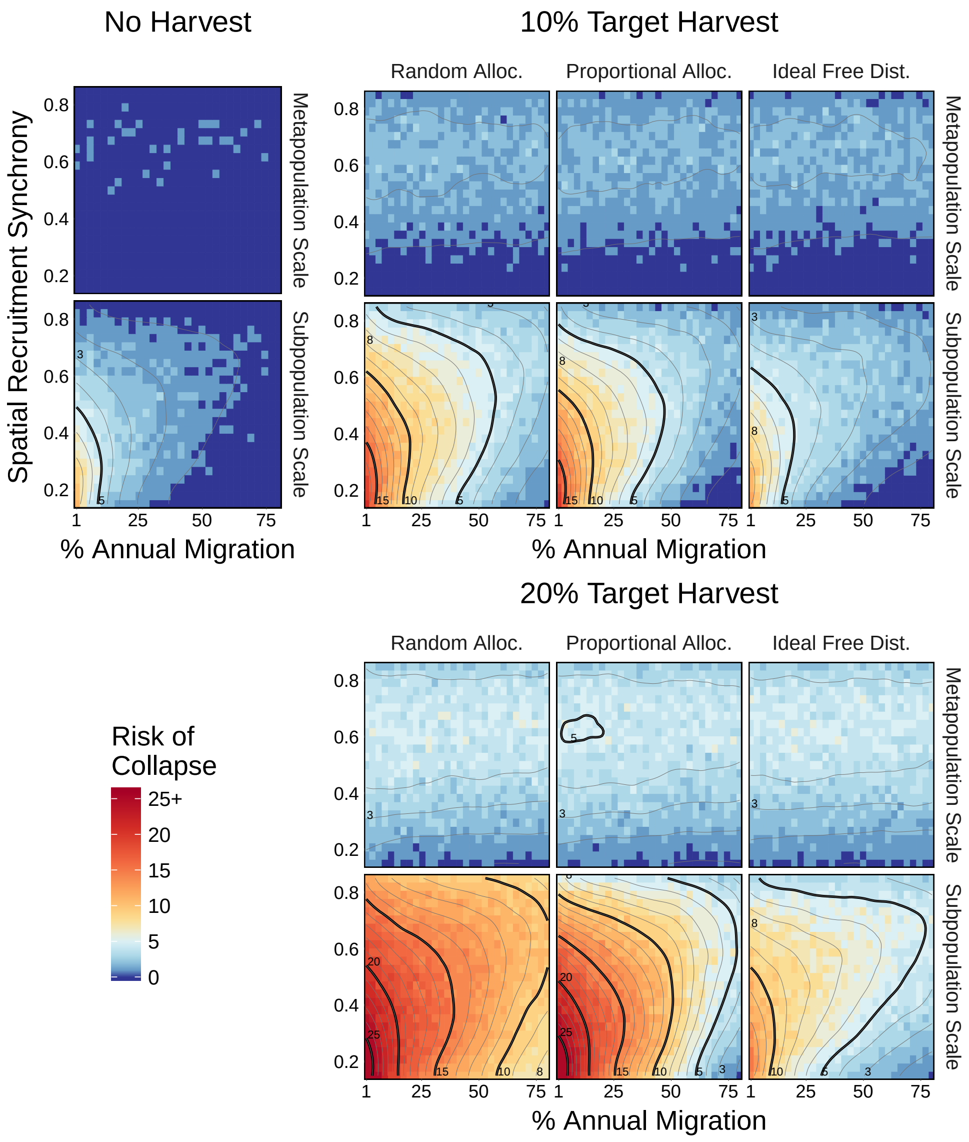


**Figure S2:** Simulation results including target harvest rates ranging from 0 to 20% for each spatial allocation strategies and measured at the metapopulation (stock) and population (substock) scales. 30% harvest rate not shown due to all risk panels above 10%. For details of allocation strategies, see *Spatial allocation of harvest* below.

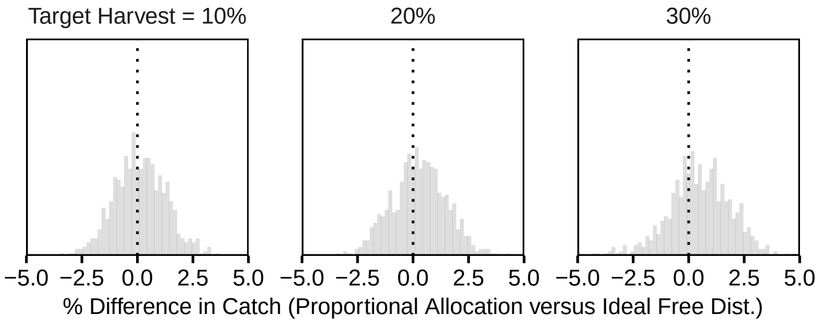


**Figure S3:** Relative frequency of the % difference in catch between proportional allocation and ideal free distribution of harvest, calculated for each individual simulation scenario as $100 * \left(\overline{catch}_{PA} - \overline{catch}_{IDF}\right) / \overline{catch}_{PA}$
.

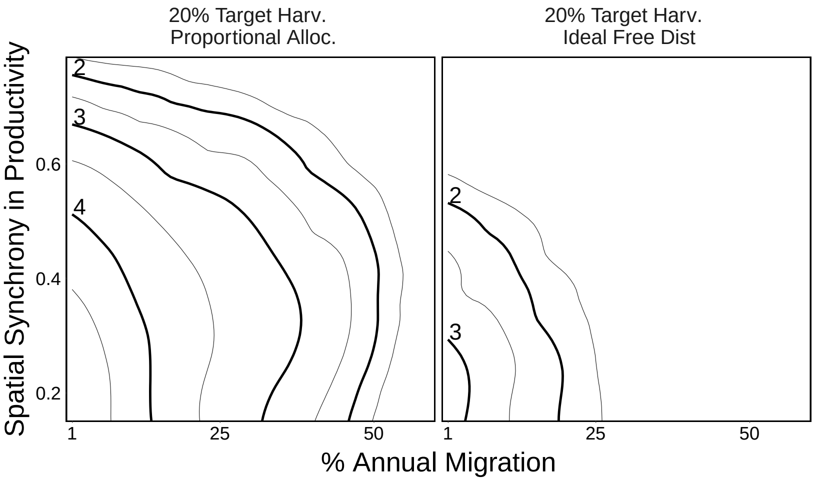


**Figure S4:** Contours representing the mean duration (years) of individual collapses under alternative spatial allocation strategies with a 20% target harvest across simulations with different simulations

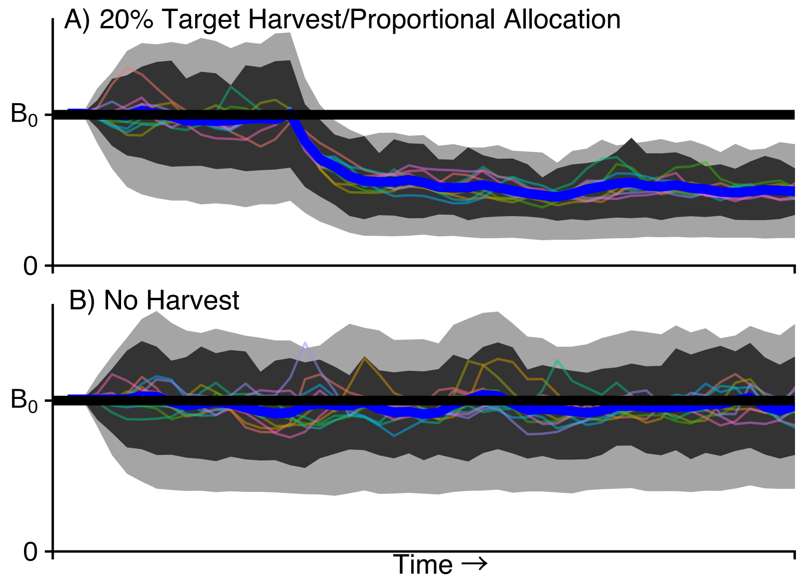


**Figure S5:** Trends from 100 simulation runs for two scenarios illustrating the average behavior of the model. The black line represents the equilibrium biomass in the absence of fishing ($B_0$); the blue line represents the mean biomass through time across all simulation runs; the dark grey band represents the 15th and 85th quantiles for the metapopulation trend across all runs; the small lines represent the mean for each subpopulation across all runs; the light grey band represents the 15th and 85th quantiles for the subpopulations across all runs

.
