## Supplementary material for "Spatial variation in exploited metapopulations obscures risk of collapse": Metadata_Appendix S4

### JOURNAL PUBLICATION CITATION:*.*Okamoto, DK, M Hessing-Lewis, JF Samhouri, AO Shelton, A Stier, PS Levin, AK Salomon. Spatial variation in exploited metapopulations obscures risk of collapse. *Ecological Applications*.

### Data Appendix S4

### Appendix S4: List of published home ranges and source information for individual studies of herring predators.

### Author(s) [of the material provided in Appendix_S4.zip]

Okamoto, Daniel K.

Florida State University

319 Stadium Drive

Tallahassee FL, 32303, USA

Hessing-Lewis, Margot

Hakai Institute

Quadra Island, Canada

Samhouri, Jameal F.

NOAA Fisheries, Northwest Fisheries Science Center

2725 Montlake Blvd E,

Seattle, WA 98112, USA

Shelton, Andrew O.

NOAA Fisheries, Northwest Fisheries Science Center

2725 Montlake Blvd E,

Seattle, WA 98112, USA

Stier, Adrian

University of California, Santa Barbara, Ecology, Evolution and Marine Biology

University of California

Santa Barbara, CA 93106, USA

Levin, Phillip S.

The Nature Conservancy

University of Washington, School of Environment and Forestry Sciences

Seattle, WA 98195, USA

Salomon, Anne K.

Simon Fraser University, School of Resource and Environmental Management

643A Science Rd,

Burnaby, BC, Canada

### File list (files found within DataS1.zip)

Appendix_S4.csv

**Description**

Appendix_S4.csv List of published home ranges and source information for individual studies of herring predators.

Number of variables: 9

Variable information by column:

1. Functional Group: Taxonomic group for species of interest
2. Species (Scientific): Genus and species name
3. Species (Common): Common name of the species
4. Range: Reported home range and units
5. Radius: Reported home range radius if reported
6. Methodology: Methodology used to assess home range
7. Paper: Name and year of the citation, all of which are cited in the main text
8. Location of study: Geographic location covered by the study
9. Notes: Comments on individual studies and caveats
